# Notch-dependent Abl signaling regulates cell motility during ommatidial rotation in *Drosophila*

**DOI:** 10.1101/2021.06.21.449310

**Authors:** Yildiz Koca, Linh T. Vuong, Jaskirat Singh, Edward Giniger, Marek Mlodzik

## Abstract

A collective cell motility event that occurs during *Drosophila* eye development, ommatidial rotation (OR), serves as a paradigm for signaling pathway-regulated directed movement of cell clusters. OR is instructed by several signaling events, including the EGFR and Notch pathways, and planar cell polarity (PCP) signaling, all of which are associated with photoreceptor R3 and R4 specification and differentiation. Here, we show that Abl kinase negatively regulates ommatidial rotation through its activity in the R3/R4 pair. Interestingly in wild-type, Abl is localized to apical junctional regions in R4 but not in R3 during OR, and this apical enrichment requires Notch signaling. We further demonstrate that Abl and Notch genetically interact during OR, and Abl co-immunoprecipitates in complexes with Notch in the developing eye disc. Perturbations of Abl interfere with adherens junction dynamics of the ommatidial preclusters, which are critical for the OR process. Taken together, our data suggest a model in which Abl kinase acts directly downstream of the Notch receptor in R4 to fine-tune OR via its input into adherens junction complexes.

## Introduction

Cells often possess directional features that play essential roles for the development, function, and homeostasis of organs and tissues. Cellular polarity across the plane of tissues, referred to as planar cell polarity (PCP), provides cells with positional information, and thus allows them to orient with respect to the body and tissue axes (Adler, 2012; Goodrich and Strutt, 2011; Humphries and Mlodzik, 2018; Peng and Axelrod, 2012). Such polarization and cellular orientation are also key for directed cellular movement within and across tissues (Butler and Wallingford, 2017; Davey and Moens, 2017; Humphries and Mlodzik, 2018). PCP has been best studied in *Drosophila* and its establishment is mediated by a specific set of evolutionarily conserved “core PCP” proteins, which include the transmembrane proteins Frizzled (Fz), Van Gogh (Vang, Vangl in vertebrates, a.k.a. Stbm in *Drosophila*) and Flamingo (Fmi, Celsr in vertebrates), and the cytoplasmic factors Dishevelled (Dsh, Dvl in vertebrates), Diego (Dgo, Diversin/Inversin in vertebrates) and Prickle (Pk) (Goodrich and Strutt, 2011; Humphries and Mlodzik, 2018; Wu and Mlodzik, 2009). During PCP establishment interactions among these core factors lead to the formation of asymmetrically localized complexes of Fz-Dsh-Dgo and Vang-Pk on opposing sides of cells, which are stabilized via intercellular homophillic adhesion of Fmi between neighboring cells across apical junctional membranes. The two complexes form separate signaling units, interacting with their set of effector proteins and thus initiate distinct tissue and cell-type specific responses (Adler, 2012; Humphries and Mlodzik, 2018; Wu and Mlodzik, 2009). Such PCP-induced downstream effector cascades can range from the (re)organization of cytoskeletal elements and the remodeling of cell adhesion complexes for the regulation of directed cell motility, to transcriptional regulation and associated cell fate changes (Adler, 2012; Davey and Moens, 2017; Devenport, 2016; Humphries and Mlodzik, 2018; Jenny, 2010). In *Drosophila*, eye development is particularly well suited to study several aspects of PCP signaling, as it entails PCP-dependent cell fate differentiation and cell motility processes (Jenny, 2010; Mlodzik, 1999; Strutt and Strutt, 1999).

The *Drosophila* eye consists of ~800 highly regularly arranged ommatidia, each of which is composed of 8 photoreceptor (R-cell) neurons (R1-R8), arranged into a defined invariant trapezoidal pattern, and 12 accessory (cone, pigment, and bristle) cells (Tomlinson and Ready, 1987; Wolff and Ready, 1991). During larval stages, the eye develops from an epithelial imaginal disc, which is initially composed of identical pluripotent precursor cells. As a wave of cell proliferation and differentiation (referred to as morphogenetic furrow, MF) travels across the disc from posterior to anterior, regularly spaced preclusters of differentiating cells start to form in its wake that will subsequently mature into ommatidial clusters (and ommatidia in the adult) (Cagan and Ready, 1989; Roignant and Treisman, 2009; Tomlinson and Ready, 1987; Wolff and Ready, 1991). At the 5-cell precluster stage, differential specification of the R3/R4 cell pair requires asymmetric Fz/PCP signaling followed by directional Notch pathway activation within the pair and EGFR-signaling, breaking the initial symmetry of the precluster (Cooper and Bray, 1999; Fanto and Mlodzik, 1999; Tomlinson and Struhl, 1999; Weber et al., 2000; Weber et al., 2008). Differential specification of R3 and R4 fates also generates the directional cues that instruct the subsequent rotation of the precluster towards the dorsal-ventral (D/V) midline, often referred to as the equator, in a process called ommatidial rotation (OR) (Blair, 1999; Jenny, 2010; Mlodzik, 1999; Strutt and Strutt, 1999). During OR, as new cells join the precluster and differentiate, the precluster collectively undergoes a 90° rotation in opposing directions in the dorsal and ventral halves of the eye and thus establish the mirror-symmetric pattern most apparent in adult ommatidia across the D/V midline (Jenny, 2010) (see also Figure 1A,C).

**Figure 1.**
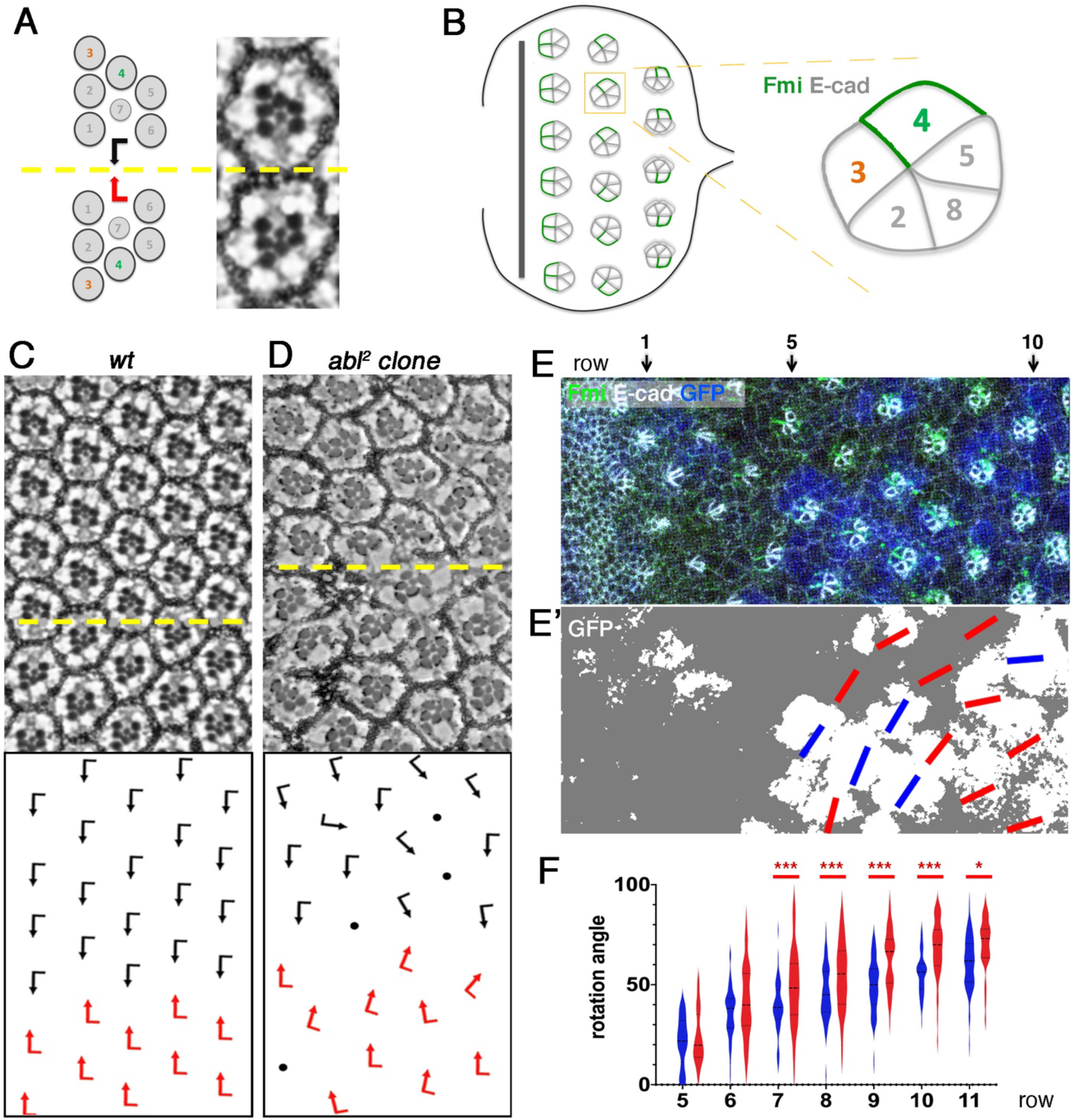
*dAbl* LOF causes various developmental defects in the eye including ommatidial misorientation. (A) Schematic and section view of the two distinct chiral forms of adult ommatidia, displaying mirror image symmetry across the equator (yellow line). (B) Schematic of third instar eye imaginal disc. As furrow (MF) moves across the eye disc from posterior to anterior, ommatidial preclusters are forming in its wake, a process that involves lateral inhibition and R8 induction. R8 subsequently induces the sequential recruitment of R2/R5 and R3/R4 precursors pairs, resulting in the 5-cell precluster. Once the symmetry of 5-cell preclusters breaks due to differential R3/R4 specification, they start to rotate towards the dorso-ventral midline until they complete a 90° rotation and are aligned perpendicular to the equator. Fmi (green), initially detected in junctions of both R3/R4 precursors, becomes enriched to R4 junctional surfaces as the precursors mature. DE-cadherin (gray) is upregulated in R2/R5 and R8 cells. Anterior is to the left and dorsal up in all panels. (C-D) Adult eye sections of wt (C) and *abl2* mosaic (D) genotypes with orientation schematics. Black and red arrows are as in A, dots indicate ommatidia with photoreceptor loss. (E) *abl2* mosaic third larval instar eye imaginal disc stained for Fmi (green), DE-cad (gray) and GFP (blue) with MF at the anterior (left). *abl2* mutant clones are marked by the absence of GFP. (E’) GFP-thresholded version of E. Wild type and mutant tissue is depicted in white and gray respectively. Accordingly, the R2/R5 plane of wild type and mutant ommatidial clusters were marked by blue and red lines respectively. (F) Quantification of rotation angles observed in individual preclusters in rows 5–11, plotted for wild type ommatidia (blue) and *abl2* mutant/mosaic ommatidia (green). Statistical analyses were performed for each row between wild type and *abl2* mosaic genotypes. Asterisks denote significance by chi-square test (\**p* < 0.05, \*\*\**p* < 0.0005). Note the over-rotation trend in the mosaic ommatidial clusters. Scale bars indicate 10 μm.

In core PCP mutants, differential R3/R4 specification fails or becomes randomized and ommatidia are often misoriented (Das et al., 2002; Wolff and Rubin, 1998; Wu et al., 2004; Zheng et al., 1995), suggesting that PCP signaling not only dictates the R3/R4 cell fate specification (Cooper and Bray, 1999; Fanto and Mlodzik, 1999), but also the direction and degree of OR. To date, several OR-specific regulators have been discovered based on the ommatidial misorientation phenotypes associated with their mutants (Brown and Freeman, 2003; Choi and Benzer, 1994; Chou and Chien, 2002; Fiehler and Wolff, 2007, 2008; Gaengel and Mlodzik, 2003; Mirkovic et al., 2011; Mirkovic and Mlodzik, 2006; Winter et al., 2001). For example, it is established that Fz/PCP signaling feeds into cadherin-based cell adhesion machinery through downstream effectors to regulate the OR process (Mirkovic et al., 2011). Furthermore, cytoskeletal reorganization of ommatidial cells is coordinated with adhesion remodeling to drive the OR process downstream of several signaling pathways, including Fz/PCP, EGFR, and Notch signaling (Brown and Freeman, 2003; Fiehler and Wolff, 2007; Gaengel and Mlodzik, 2003; Koca et al., 2019; Winter et al., 2001). Despite this knowledge, actual mechanistic insights into the OR process remain largely elusive.

Abelson (Abl) kinases are a family of non-receptor tyrosine kinases that govern a multitude of cellular processes in metazoans, including proliferation, differentiation, survival, and migration (Bradley and Koleske, 2009; Hernández et al., 2004; Wang, 2014). Unlike master switch kinases, Abl kinases are transitionally activated and/or subcellularly enriched, depending on the function they are involved in and can therefore regulate distinct cellular processes (Bradley and Koleske, 2009; Hernández et al., 2004; Wang, 2014). In the absence of activating signals, the SH2 and SH3 domains of Abl interact with each other and lock the protein in a kinase-inactive state (Hantschel et al., 2003; Nagar et al., 2003). Although a direct mechanism for Abl kinase activation has not been identified, it is likely that upon stimulatory signals, the SH2 and SH3 domains interact locally with secondary molecules, unlocking the inhibitory conformation and locally enabling kinase activity. Remarkably, Abl kinases can be activated downstream of various signals, including growth factors, cell-ECM interactions, or adhesion receptors, highlighting the versatile nature of Abl signaling in cells (Bradley and Koleske, 2009; Hernández et al., 2004; Wang, 2014). *Drosophila* Abl (dAbl) is primarily cytoplasmic and has been linked to the regulation of cytoskeletal and adhesion processes in a number of tissues. During early embryo morphogenesis, for example, dAbl is required to localize actin polymerization and actomyosin activity to the apical domain by restricting Enabled (Ena) activity (Baum and Perrimon, 2001; Fox and Peifer, 2007; Grevengoed et al., 2003; Grevengoed et al., 2001; Jodoin and Martin, 2016). In axon guidance, dAbl feeds into multiple signaling branches, including Ena and Rac GTPases, to regulate the balance of linear versus branched actin networks (Kannan et al., 2017; Song et al., 2010). Germ band elongation requires dAbl to locally control the mobility of adhesion complexes through phosphorylation of Arm/β-cat thus promoting convergent extension movements (Tamada et al., 2012). In addition, during photoreceptor morphogenesis dAbl has been shown to be essential for the maintenance of the apicobasal integrity in photoreceptors and associated proper organization of adherens junctions (Xiong and Rebay, 2011). Interestingly, dAbl was also shown to function within the core PCP pathway, through Dsh phosphorylation, in order to promote Fz/Dsh-PCP signaling within R3/R4 pairs (Singh et al., 2010). The involvement of Abl kinases in cell adhesion and cytoskeletal remodeling has also been documented in various vertebrate contexts, suggesting that many of its functions are conserved across species (Bradley and Koleske, 2009; Hernández et al., 2004; Wang, 2014).

Here we show that during *Drosophila* eye patterning dAbl kinase negatively regulates ommatidial rotation downstream of the Notch receptor. Loss and gain-of-function genotypes of dAbl consistently cause opposite effects on the OR process. dAbl becomes apically enriched in photoreceptors R8, R2/R5 and importantly R4 but not in R3 during OR. The apical junctional enrichment in R4 requires Notch signaling. Functionally, dAbl and Notch interact genetically during OR, and dAbl co-exists in complexes with Notch in the developing eye disc. Our data collectively suggest that Abl acts directly downstream of the Notch receptor in R4 by acting on the cadherin/β-catenin complexes and cell adhesion, and thus it serves a “brake function” during the OR process, fine-tuning the OR cell motility for convergence to a 90° angle.

## Results

### Abl kinase regulates ommatidial rotation

Our studies in the past suggested that dAbl could have a role in the ommatidial rotation (OR) process, as overexpression of the kinase caused OR defects, but this potential dAbl contribution to *Drosophila* eye patterning and morphogenesis remained largely unexplored (Singh et al., 2010). To investigate how dAbl contributes to OR during retinal patterning, we analyzed the phenotypes of loss-of-function (LOF) clones of *dAbl* in mosaic eye tissue. While clones of the *abl^2^* allele can be recovered in adult eyes, they display pleiotropic developmental and morphogenetic defects (Figure 1C-D). Besides the previously reported phenotypes of loss and malformation of photoreceptors (Singh et al., 2010; Xiong and Rebay, 2011) and loss of chirality (Singh et al., 2010), these clones also display misoriented ommatidia, consistent with the hypothesis that dAbl also regulate the OR process (Figure 1C-D). To establish that the OR defects observed in LOF clones are a primary phenotype of *dAbl* mutant clusters, we examined *abl^2^* clonal tissue in eye imaginal discs at the time of OR. *dAbl* mutant or mosaic ommatidial clusters frequently displayed an over-rotation phenotype in rows 5-11 as compared to wild-type clusters (Figure 1E-F). In such early *dAbl* deficient ommatidial (pre)clusters, the organization of the 5-cell precluster was largely normal (Figure 1E-F), suggesting that Abl kinase has an OR-specific function during eye development.

To get further insight into the role of dAbl in OR, we analyzed the effects of Abl gain-of-function (GOF) posterior to the MF by employing the *sevGal4* driver (see Suppl. Figure S2 for expression pattern). In *sevGal4*, *UAS-dAbl* adult eyes (*sev*>*Abl*), most aspects of eye development were normal, including a correct chiral ommatidial arrangement, with the exception of many ommatidia being misoriented, and notably an under-rotation pattern of ommatidia was most frequently observed (Figure 2A-B). To confirm that the OR defects observed in adult eyes upon *sev*>*Abl* expression arise as primary defects during the OR process, we analyzed the respective larval eye discs during ommatidial patterning. Consistently, dAbl overexpressing ommatidia showed markedly slower rate of rotation, as compared to wild type (Figure 2C-E). These data, together with the LOF clonal phenotypes displaying over-rotation features, suggest that Abl regulates the rate of rotation and that it has a specific inhibitory or ‘braking’ function in OR.

**Figure 2.**
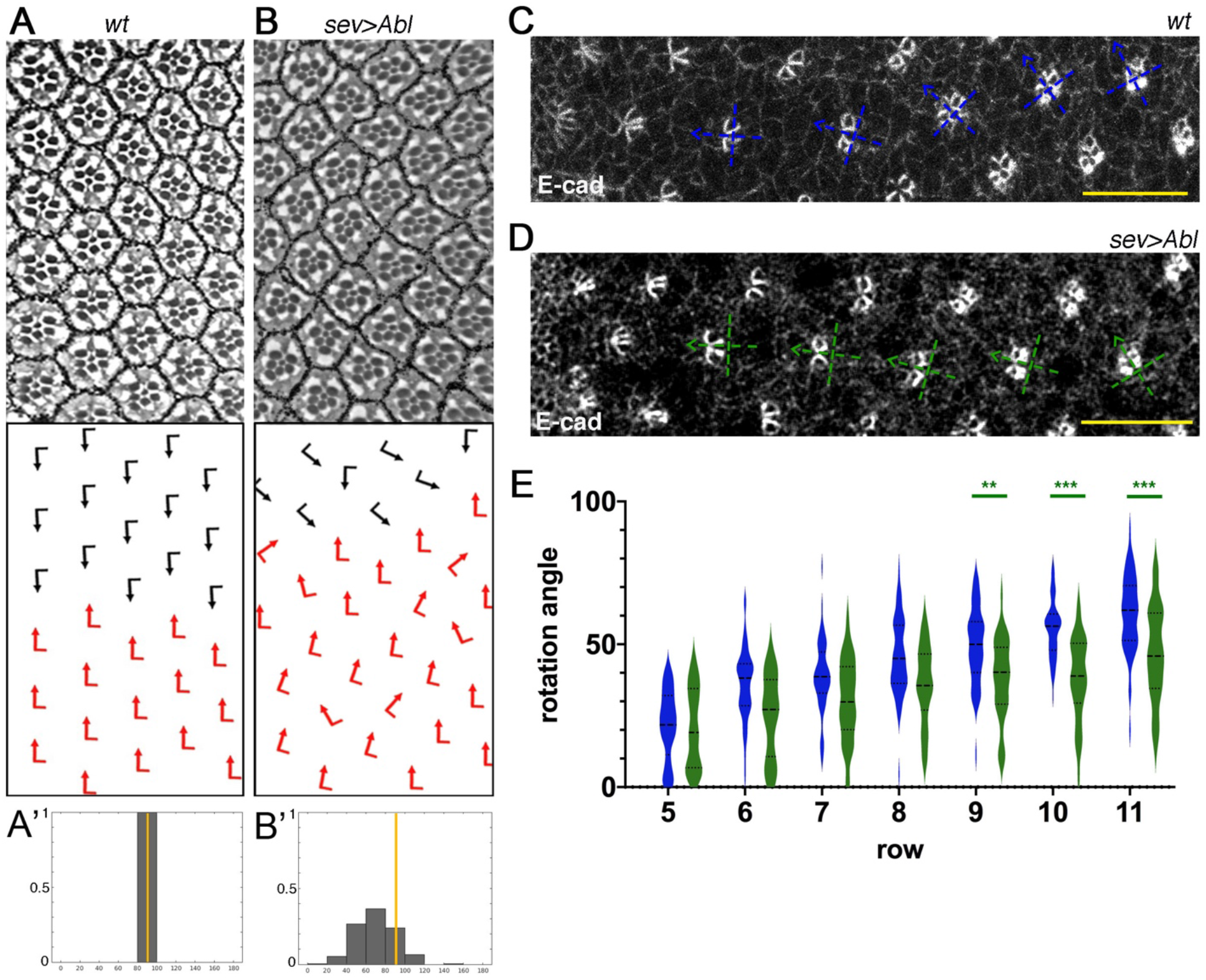
Abl overexpression in eye discs posterior to MF slows down OR. (A-B’) Adult eye sections with ommatidial orientation schematics (arrows as in Fig. 1) and orientation angle histograms of eyes of the genotypes indicated: (A,A’) *wt*; (B, B’) *sev*>*Abl*. (C-D) Third instar larval eye imaginal discs stained for DE-cad (gray) in wild type (C) and *sev*>*Abl* (D). Blue and green dashed cross-arrows, respectively, indicate the orientation of ommatidial preclusters for each genotype. (E) Quantification of rotation angles observed in individual preclusters in rows 5–11, plotted for *wt* (blue) and *sev*>*Abl* (green). Statistical analyses were performed for each row between wt (blue) and Abl OE (green) genotypes. Asterisks denote significance by chi-square test (\*\**p* < 0.005, \*\*\**p* < 0.0005). Note the strong under-rotation trend of ommatidia in multiple rows of *sev*>*Abl* genotype compared to wild type. Scale bars indicate 10 μm.

### Abl kinase is apically enriched and required in R4 to slow down OR

To gain insight into how dAbl may regulate rotation, we first analyzed the expression and localization pattern of dAbl during OR in eye imaginal discs. dAbl expression was detected prominently in photoreceptors starting from a few rows posterior to the MF and persisting throughout ommatidial development. Notably, dAbl became enriched apically at the level of the adherens junctions in R8, R2/R5 and R4 shortly after the initial fast phase of OR, a pattern maintained as the preclusters matured (from row 8 onward; Figure 3A-C’). Interestingly, we did not observe apical localization of dAbl in R3 at any stage posterior to the MF. Thus, dAbl displayed a striking localization difference in the apical junctional area between the two cells of the R3/R4 pair (summarized in Figure 3B’’). Unlike this apical pattern, Abl localization and intensity did not show a notable difference between R3 and R4 more basally (Suppl. Figure S3), suggesting that Abl is specifically enriched in the apical domain of R4 but not R3 during the OR process.

**Figure 3.**
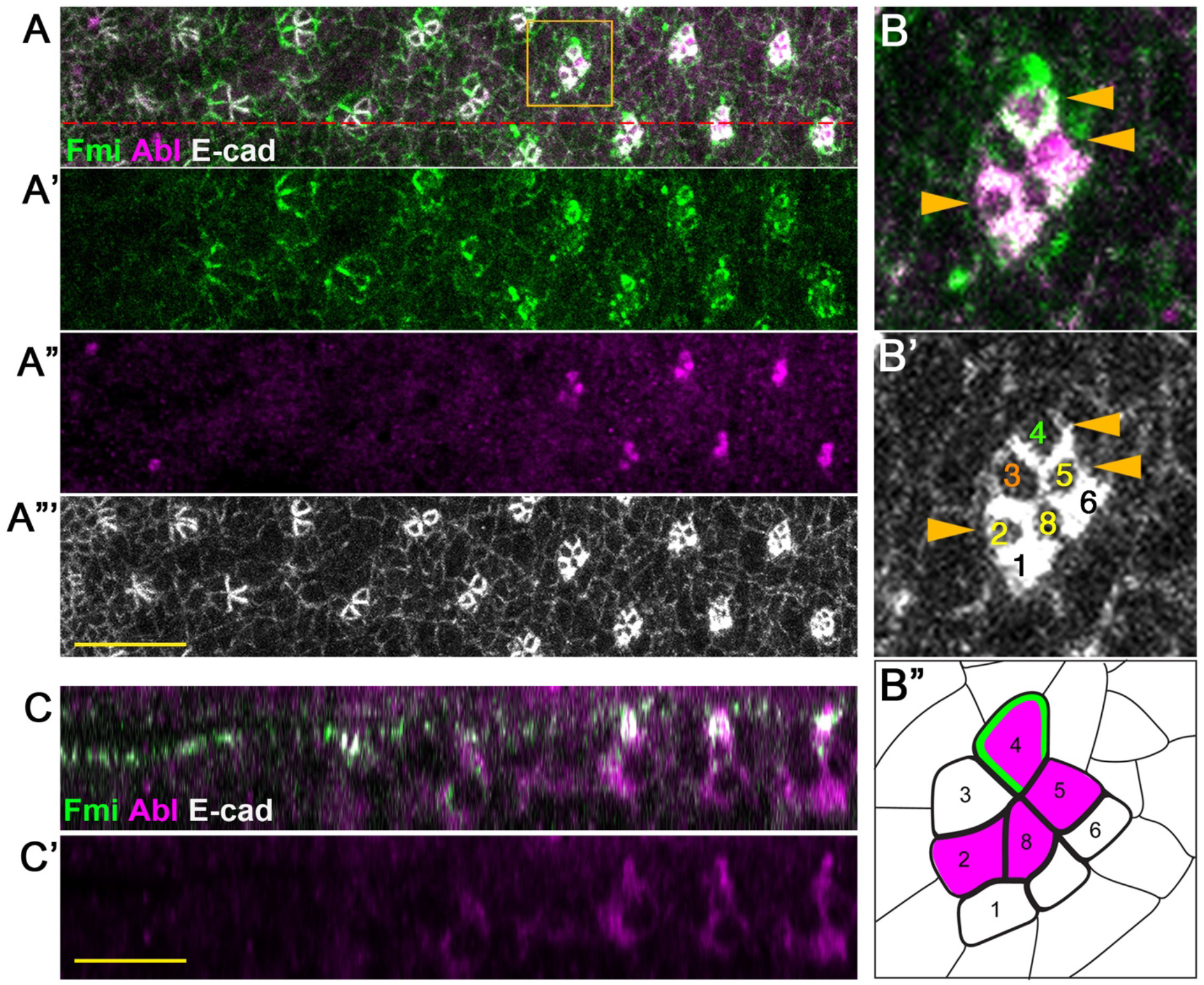
Abl is apically enriched in all photoreceptors of the 5-cell precluster but R3. (A-A’’’) Control third instar larval eye imaginal disc stained for Fmi (green), Abl (magenta) and DE-cad (gray). (B-B’’’) Zoomed view of the individual cluster boxed in A (B), monochrome of E-cad to highlight indivudal cells (B’), and schematized (B’’). Note that Abl is present in apical junctional regions in R8, R2/R5, and R4, but not in R3. (C-C’) xz-view of the eye disc staining, at the section marked by the red dashed line in A. Note the localization of Abl at the level of apical DE-cad and Fmi. Scale bars indicate 10 μm.

Differential specification of R3/R4 by Fz/PCP and Notch signaling and the associated signaling events are critical for the correct direction and execution of OR (Blair, 1999; Mlodzik, 1999; Strutt and Strutt, 1999). To understand whether the contrasting apical localization pattern of dAbl between R3/R4 is important for OR, we aimed to perturb dAbl function and localization specifically in cells of the R3/R4 pair. To this end, we employed the *mδ0.5-Gal4* driver which is initially active in both R3/R4 precursors and later becomes specifically upregulated in R4 as a result of Notch-mediated R4 specification (Cooper and Bray, 1999; Koca et al., 2019). *mδ0.5-Gal4* mediated knock-down (KD) of *dAbl*, within R3/R4 pairs led to a significant aberration in the rotation pattern of ommatidial clusters as compared to wild-type, with clusters generally rotating faster (Figure 4A-B,D; Suppl. Figure S4). Conversely, dAbl over-expression in R3/R4 with *mδ0.5-Gal4* caused a significant under-rotation pattern starting from early stages of OR (Figure 4A,C-D; and also Suppl. Figure S4). Taken together, these data suggest that the differential dAbl localization pattern, with dAbl being highly enriched apically in R4, is required to fine-tune the slowing rate of rotation at later stages of the process.

**Figure 4.**
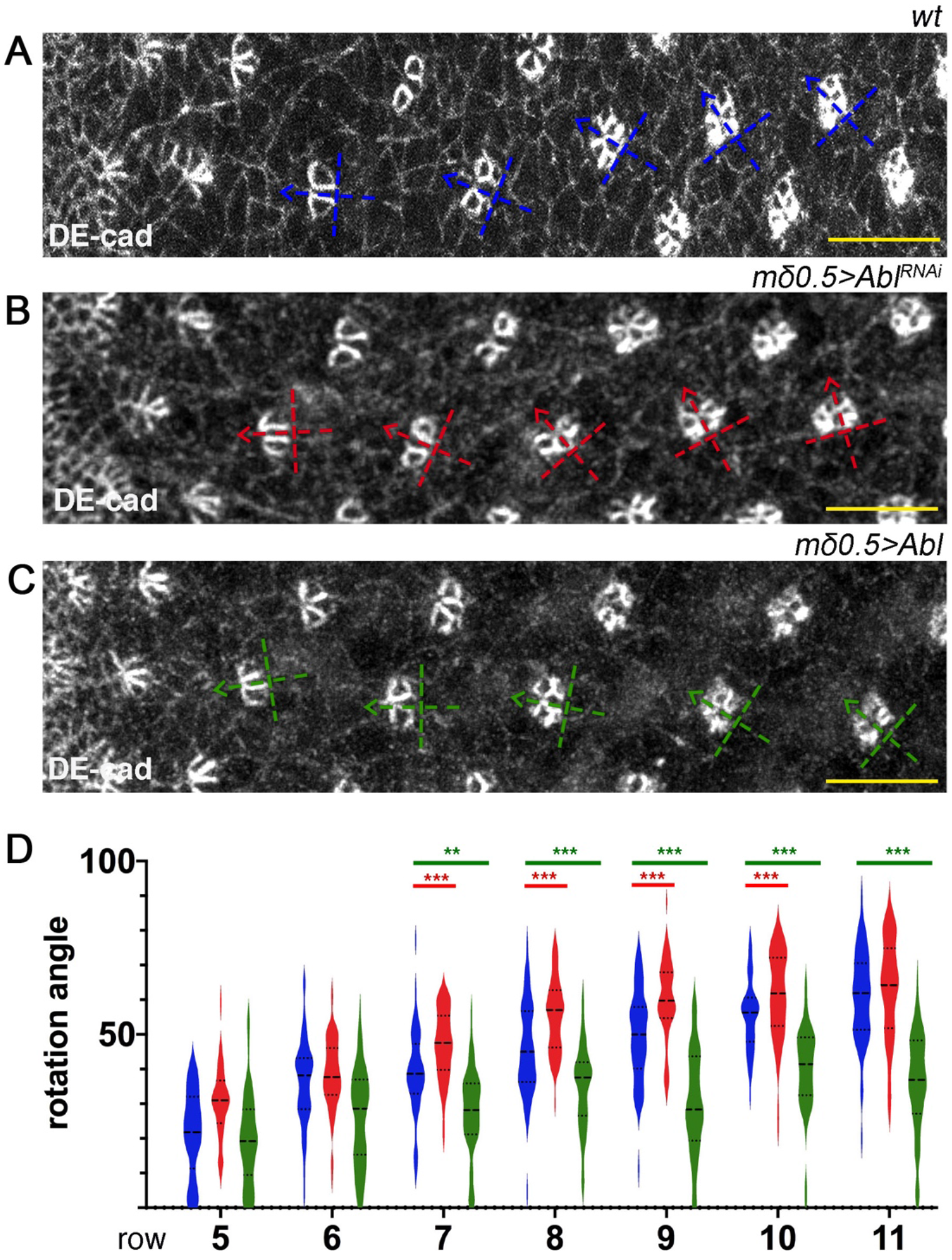
Abl signaling within R3/R4 pair negatively regulates OR. (A-C) Third instar larval eye imaginal discs stained for DE-cad (gray) in wt (A), *mδ0.5*>*Abl^RNAi^* (B) and *mδ0.5*>*Abl* (C). Blue, red and green dashed cross-arrows, respectively, indicate the orientation of ommatidial preclusters for each genotype. (D) Quantification of rotation angles observed in individual preclusters in rows 5–11, plotted for wt (blue), *mδ0.5*>*Abl^RNAi^* (red) and *mδ0.5*>*Abl* (green). Statistical analyses were performed for each row between wt (blue) and Abl RNAi (red) or Abl OE (green) genotypes. Asterisks denote significance by chi-square test (\*\**p* < 0.005, \*\*\**p* < 0.0005). Note the over- and under-rotation trends observed in Abl RNAi (red) or Abl OE (green) genotypes, respectively, relative to wt. Scale bars indicate 10 μm.

### Notch activation is required for apical enrichment of dAbl in R4

The stark difference in apical localization of dAbl between the R3 and R4 cells raised the possibility that either Fz/PCP or Notch-signaling, both required to specify R4, instruct apical dAbl localization. This possibility was investigated in the R3/R4 cell pair both in core PCP mutant eye clones and upon Notch-signaling interference. Firstly, we did not observe any difference in apical dAbl localization pattern of R3/R4 pairs in *pk*^−/−^, *stbm*^−/−^ double mutant clones, as compared to wild type (Figure 5A-A’). Among the core Fz/PCP signaling genes, both *pk* and *stbm/Vang* are functionally required in R4 (Jenny et al., 2003; Wolff and Rubin, 1998), therefore these data suggest that Fz/PCP-signaling is not directly required for apical dAbl localization in R4, and its associated differential localization pattern between cells of the R3/R4 pair. Nevertheless, dAbl appears to interact with core PCP factors to regulate OR. For example, while over-expression of Fmi in R3/R4 pairs with the *mδ0.5-Gal4* driver leads to the randomization of chirality, it only occasionally leads to misorientation in the adults; however, when Fmi and dAbl were co-over-expressed (*mδ0.5*>*Fmi*, >*Abl*), the misorientation phenotype was exacerbated in comparison to the individual backgrounds (*mδ0.5*>Abl and *mδ0.5*>Fmi; Suppl. Figure S5A-D, see figure legend for more details).

**Figure 5.**
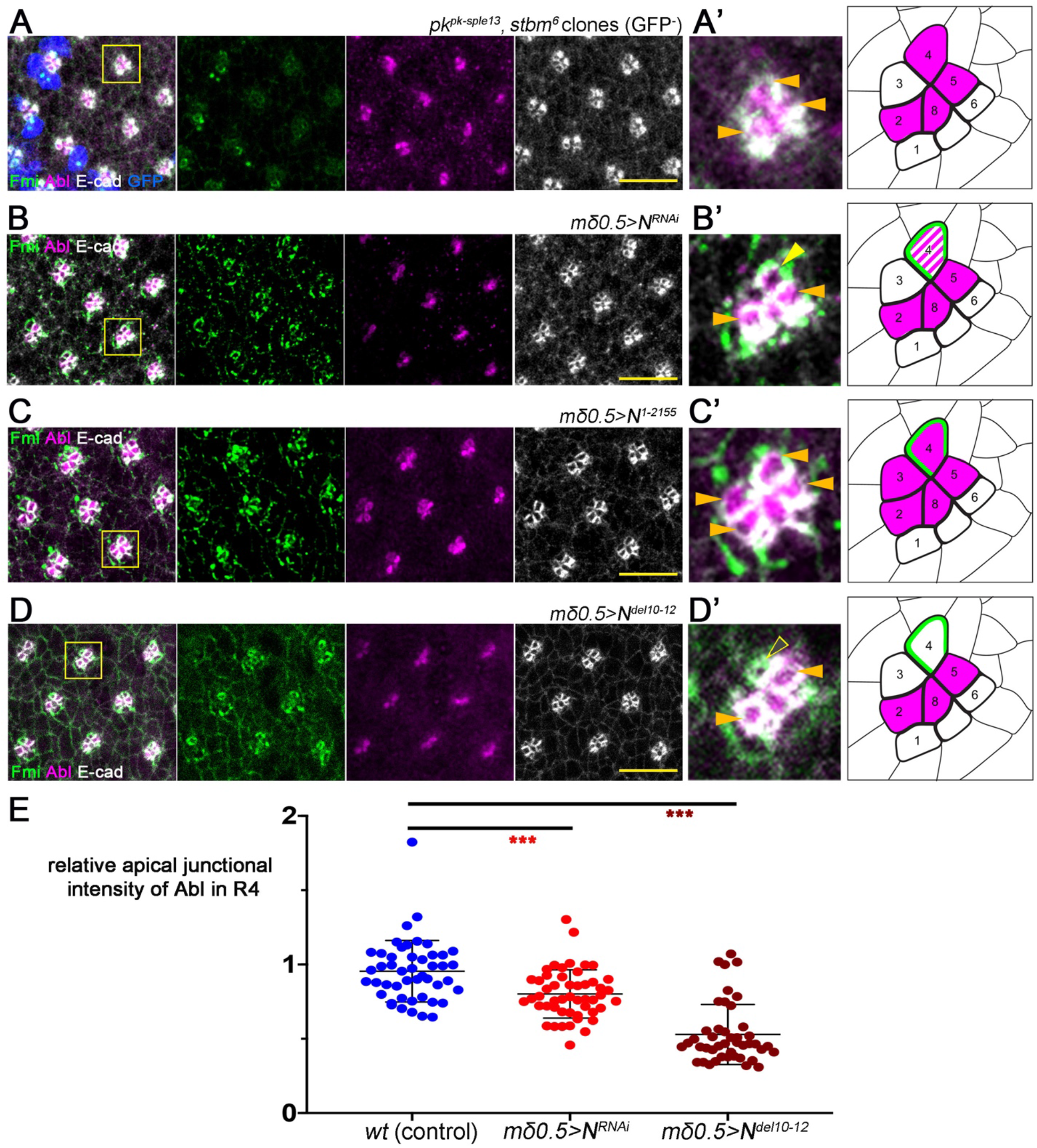
Notch signaling, but not core PCP factors, is required for apical Abl localization in R4. (A-D) Third instar larval eye imaginal discs of the designated genotypes stained for Fmi (green), Abl (magenta) and DE-cad (gray): (A) *pk^pk-sple13^stbm^6^* double mutant mosaics, mutant clones are marked by the absence of GFP (blue), (B) *mδ0.5*>*N^RNAi^*, (C) *mδ0.5*>*N^1-2155^*, (D) *mδ0.5*>*N^del10-12^.* (A’-D’) High magnification individual clusters for each genotype (marked by yellow boxes in A-D) ar depicted in left panels, depicted in right panels are the respective schematics (same schematic drawing as in Fig 3B’). Note that apical junctional enrichment in R4 is not affected in *pk, Vang* double mutant clones, but is reduced or lost in genotypes affecting Notch signaling (*mδ0.5* > *N^RNAi^* and *mδ0.5* > *N^del10-12^* backgrounds, B’ and D’, respectively); in contrast Notch activation in R3 recruits Abl to junctional apical regions in this cell as well (C’). (E) Quantification of apical Abl intensity in R4s, normalized to that in R2, plotted for individual clusters in *wt* (blue), *mδ0.5* > *N^RNAi^* (red), and *mδ0.5* > *N^del10-12^* (brown). Asterisks denote significance by chi-square test (\*\*\**p* < 0.0005). Note the normalized reduction of apical Abl intensity in R4 in the *mδ0.5* > *N^RNAi^* and *mδ0.5* > *N^del10-12^* backgrounds, as compared to *wt*. Scale bars indicate 10 μm.

Notch is specifically and asymmetrically activated in R4, as a result of directional Fz/PCP-signaling between R3/R4, and thus we next asked whether Notch signaling was required for apical dAbl enrichment in R4. Strikingly, apical junctional localization of dAbl was largely diminished in R4 upon Notch knockdown (via *mδ0.5-Gal4*) (Figure 5B-B’ and E), while the apical, junctional R4 marker Fmi remained intact. Furthermore, when a stable, truncated isoform of Notch (which behaves like an activated isoform; Suppl Fig. S5F) was expressed in the R3/R4 pair (*mδ0.5-Gal4, UAS-N^1-2155^)*, dAbl was apically localized in both cells of the pair (Figure 5C-C’). These data are consistent with the hypothesis that (activated) Notch is sufficient for apical accumulation of dAbl, which in wild-type is restricted to R4. Hence, we hypothesize that apical accumulation, specific to R4 (but not to R3), requires Notch activation.

To further elucidate how Notch mediates apical dAbl localization and whether it requires ligand-mediated activation and Notch pathway signaling down to its transcriptional activity, we have analyzed several truncated versions of Notch (expressed under the *mδ0.5-Gal4* driver) and investigated if and how these could affect apical recruitment of dAbl in R4. Strikingly, expression of a truncated form of Notch that cannot bind to its ligand Delta, termed *N^del10-12^* (lacking the EGF repeats 10-12 in its extracellular ligand binding domain, which are required for its interaction with Delta (Le Gall et al., 2008; Rebay et al., 1991)), displayed a marked loss of apical localization of dAbl in R4 (Figure 5D, D’ and E; while the localization of the R4 apical marker Fmi appeared normal as a control). Notably, in this background, the preceding Notch-signaling mediated R4 fate specification was unaffected and the Notch transcriptional reporter, *mδ0.5-lacZ*, was still active in R4 cells of the rotating ommatidia (Suppl Fig. S5E), suggesting that Delta binding to Notch is required for apical dAbl localization potentially via a transcription-independent mechanism.

### dAbl associates with a junctional Notch complex

We next asked whether dAbl can physically associate with Notch *in vivo*. Strikingly, Notch co-immunoprecipitated with dAbl in extracts from third instar larval eye discs (Figure 6A-B). Consistently, Notch also co-immunoprecipitated with the adherens junction component Arm/β-catenin (which is a known Abl phosphorylation target; Tamada et al., 2012) in eye disc tissue, suggesting that Notch interacts with adherens junction components. Nevertheless, Notch and dAbl did not interact directly in a GST-pulldown assay (Figure 6C), indicating that Notch and dAbl co-exist in membrane associated junctional complexes during eye development, but that additional factors are required to link the two proteins. To further assess whether Notch can generally instruct the localization of dAbl to junctional membrane regions, we turned to a salivary glands assay, where cells are large in size and cell compartments are relatively easy to analyze. In wild-type salivary glands, dAbl is detected in the cytoplasm and at junctional membranes, where it co-localizes with E-cad and Notch (Figure 6F). Importantly, junctional dAbl localization was increased in salivary gland cells upon Notch over-expression (Figure 6G-H, also Suppl. Fig. S6), while total dAbl levels were unchanged in this background, suggesting that the Notch receptor can generally potentiate dAbl localization to adherens junctions. Taken together, these results argue that the Notch receptor promotes the recruitment of dAbl to apical junctional complexes.

**Figure 6.**
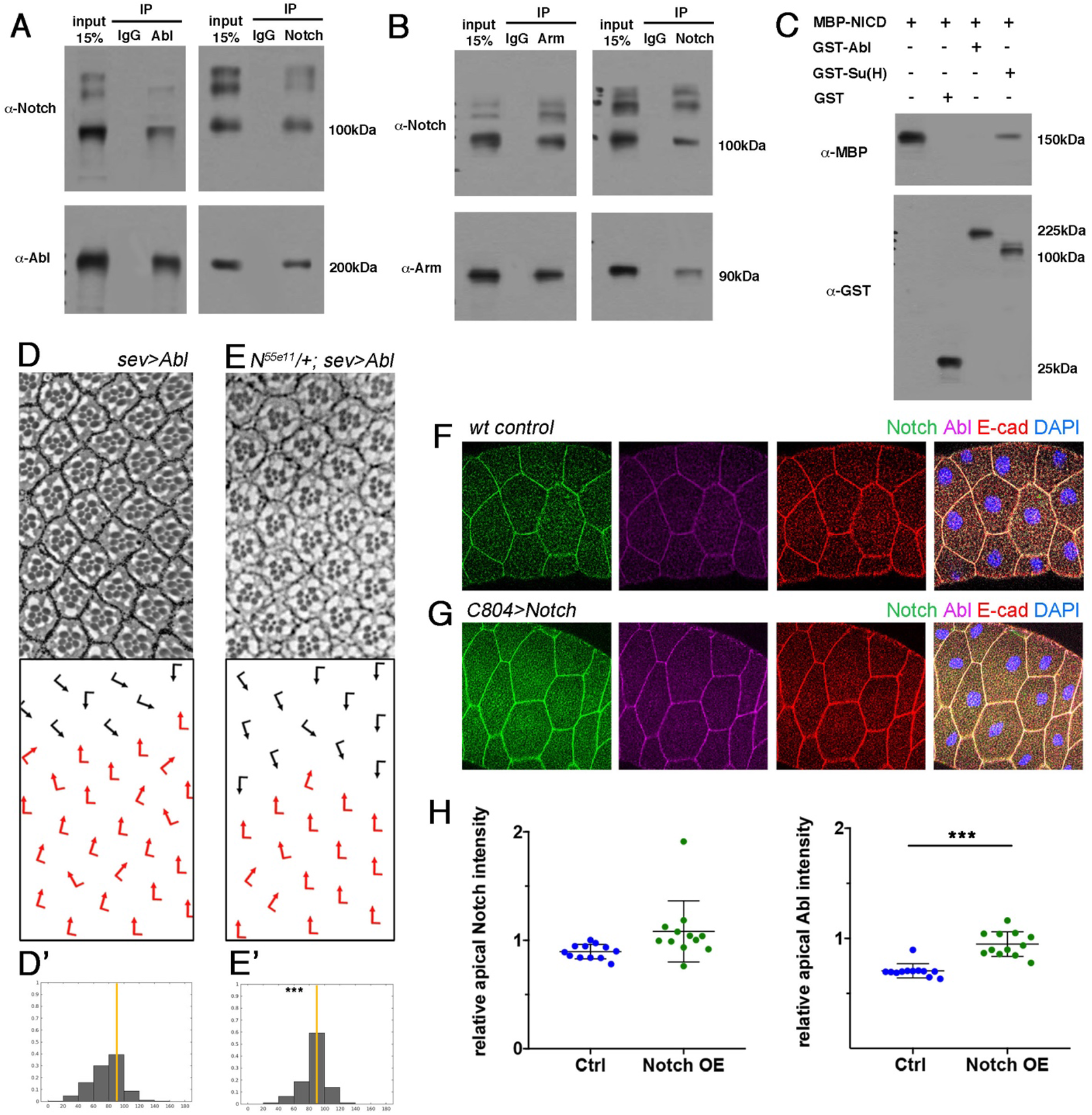
Abl and Notch physically and genetically interact to regulate OR. (A) Wild type third larval instar eye disc lysates immunoprecipitated by α-IgG, α-Abl or α-Notch, blotted for Abl and Notch. Note that Abl and Notch immunoprecipitate with each other. (B) Wild type third larval instar eye disc lysates immunoprecipitated by α-IgG, α-Arm or α-Notch, blotted for Arm and Notch. Note that Arm and Notch immunoprecipitated with each other. (C) MBP-NICD pulled down by GST-Su(H) but not GST-Abl or GST. (D-E’) Adult eye sections with ommatidial orientation schematics and orientation angle histograms of eyes of the genotypes indicated: (D,D’) *sev*>*Abl*, (E, E’) *N^55e11^*/+; *sev*>*Abl*. Asterisks denote significance by chi-square test (\*\*\**p* < 0.0005). (F-G) *wt* (F) and Notch overexpressing (G) salivary gland cells stained for Notch (green), Abl (magenta), E-cad (red), and DAPI (nuclei in blue). Note the enrichment of Abl at the apical junctions upon Notch overexpression. (H) Quantification of junctional Abl and Notch intensities relative to junctional E-cad (as detected from equivalent samples to those shown F-G). Note marked increase in junctional Abl levels upon Notch overexpression (right graph). Asterisks denote significance by both chi-square test and Student’s t-test (\*\*\**p* < 0.0005).

To confirm that the association of Notch and dAbl is important for the function of dAbl during OR, we tested for genetic interactions between Notch and dAbl specifically in the OR context. To this end, we used the OR-specific dAbl GOF genotype, *sev*>*Abl*, which displays under-rotation of ommatidial clusters (see above, Figure 2). Strikingly, removing a gene copy of *Notch* (and thus reducing Notch protein levels) we observed a marked suppression of the *sev*>*Abl* misrotation phenotype in the adults (Figure 6D-E’). This observation is consistent with the notion that dAbl acts downstream of the Notch receptor and requires Notch for its localization and/or function to regulate the OR process.

### dAbl interacts with junctional components during OR

Abl kinases mediate cytoskeletal and junctional dynamics in various contexts (Bradley and Koleske, 2009; Hernández et al., 2004; Wang, 2014). Adherens junction components have been shown to be critical for the OR process (Mirkovic et al., 2011; Mirkovic and Mlodzik, 2006). As dAbl is enriched in the apical junctional region of R4, we asked whether dAbl affects junctional components in the OR context. As the Abl GOF genotypes (e.g. *sev*>*Abl* or *mδ0.5*>*Abl*, see above) caused (largely) clean OR defects, we used a clonal GOF strategy to examine dAbl effects on junctional components in mosaic third larval instar eye discs. During OR, N-cadherin (N-cad) becomes enriched at the junctional border between the R3/R4 cells (Mirkovic and Mlodzik, 2006). Strikingly, clonal overexpression of dAbl in R3/R4 cells led to a reduction in N-cad levels at the apical junctional R3/R4 borders (Figure 7A-A’), suggesting that dAbl regulates the R3/R4 adherens junction dynamics. In contrast, comparing the apical F-actin pattern of R3/R4 cells between dAbl GOF and wild-type eye discs, we did not detect apparent differences, with F-actin being densely localized at the apical junctional ring of each photoreceptor in both genetic backgrounds (Figure S7).

**Figure 7.**
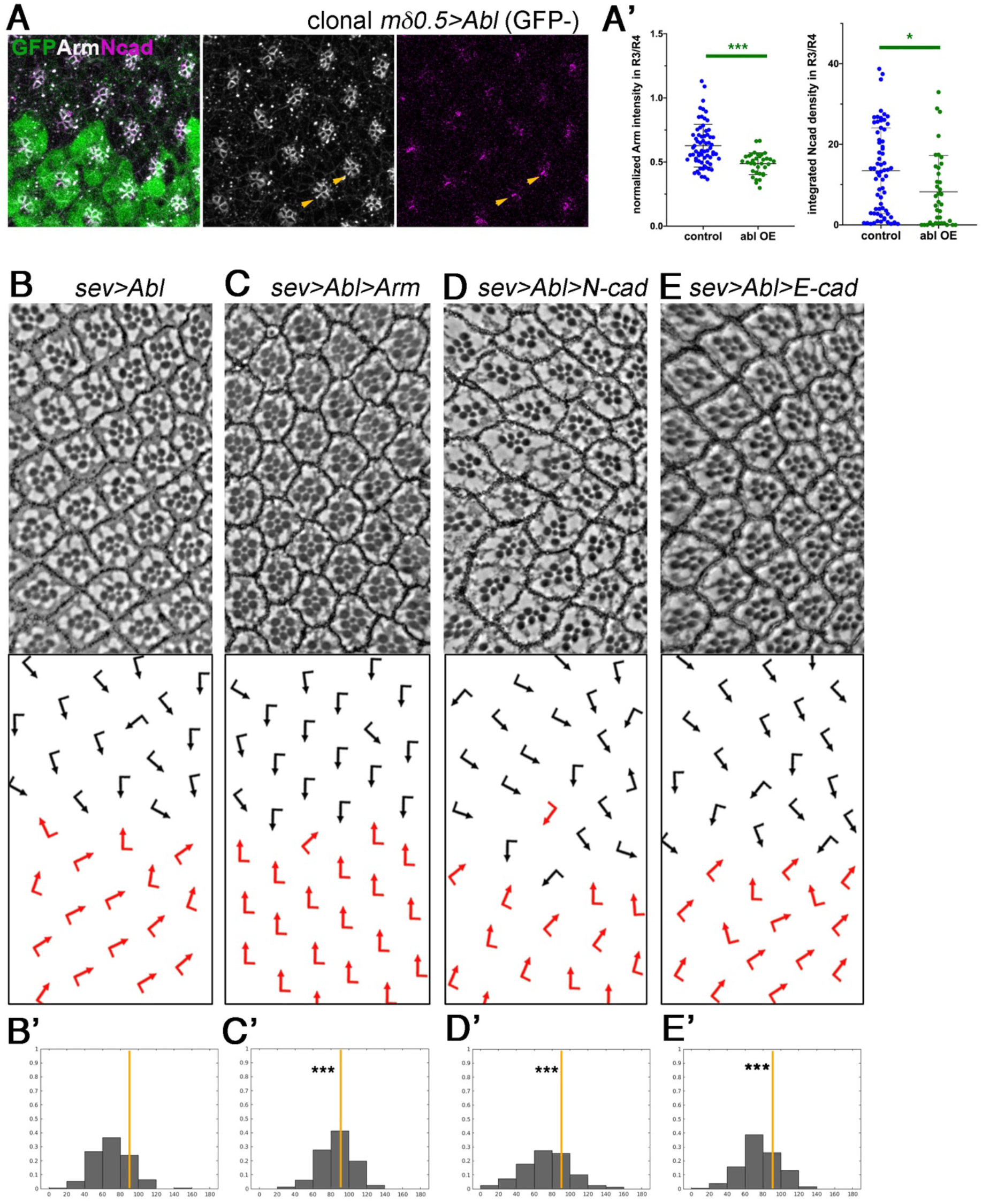
Abl feeds into junctional dynamics to regulate OR. A) *mδ0.5*>*Abl* mosaic eye disc stained for GFP (green), N-cad (magenta) and Arm (gray). *mδ0.5*>*Abl* tissue is marked by the absence of GFP. A’) Quantification of integrated apical N-cad intensity and normalized Arm intensity within the R3/R4 border, plotted for individual clusters in wt (blue) and *mδ0.5*>*Abl* (green). B-E’) Adult eye sections with ommatidial orientation schematics and orientation angle histograms of eyes of the genotypes indicated: B,B’) *sev*>*Abl*, C,C’) *sev*>*Ab*>*Arm*, D,D’) *sev*>*Abl*>*DN-cad* E,E’) *sev*>*Abl*>*DE-cad*. Asterisks denote significance by chi-square test (\**p* < 0.005, \*\*\**p* < 0.0005).

To functionally confirm the effect of dAbl on adherens junctions during OR, we used again the rotation specific dAbl GOF scenario (in *sev*>*Abl*). Strikingly, *sev*>*Abl* eyes displayed markedly suppressed under-rotation features when dAbl was co-over-expressed with the junctional component β-catenin (Armadillo/Arm in *Drosophila*; *sev*>*Abl*, >*Arm*) (Fig. 7B-C). Note that *sev*>*Abl* associated under-rotation phenotype (with many clusters rotating significantly less than 90°; Figure 7B-B’) was largely suppressed and an average rotation of close to 90° was restored in the *sev*>*Abl*, >*Arm* background (Figure 7C-C’). These data are consistent with the notion that dAbl regulates OR via its effects on adherens junctions. As co-over-expression of *Drosophila* E-cad or N-cad did not suppress the *sev*>*Abl* phenotype (Figure 7D-E), it appeared that Arm/β-catenin might be a more direct target of Abl regulation, which is consistent with dAbl phosphoprylating Arm (Tamada et al., 2012) (see Discussion).

## Discussion

Abl has been shown to regulate multiple aspects of cell migration in various contexts from *Drosophila* to vertebrates. Our results argue that dAbl regulates cell motility during OR. Although loss of Abl function interferes with multiple aspects of photoreceptor development and morphogenesis, including R3/R4 chirality and ommatidial orientation, overexpression of dAbl in developing ommatidial clusters in the eye disc affects quite specifically OR, suggesting that dAbl has an OR-specific function. In the OR context, dAbl appears to have an inhibitory role, as ommatidial clusters with increased dAbl levels under-rotate, whereas *dAbl* mutant/mosaic ommatidia tend to rotate faster.

The localization pattern of dAbl posterior to the MF provides further insight about its role in OR. dAbl becomes apically enriched in photoreceptors R8, R2/R5 and R4, following a steady phase of rotation, at the time when clusters start to slow down and refine their motility until the completion of the 90° angle. The under-rotation features observed upon dAbl overexpression is consistent with the notion that dAbl becomes apically localized in specific R-cells, towards the later stages of OR, to slow down the process. Interestingly, there is a differential localization pattern between R3 and R4 in terms of dAbl apical junctional association. Considering the role of the R3/R4 pair and associated signaling pathways in OR, it is tempting to speculate that the differential dAbl localization in R3 and R4 is comparable to the requirement of the Nmo kinase within R3/R4, with Nmo providing a directional impulse to rotation in R4 (Mirkovic et al., 2011) and dAbl regulating its slowing down. Our data argue that dAbl activity within R3/R4 pairs is indeed important for fine-tuning OR. Knockdown and overexpression of dAbl, specifically in R3/R4 pairs, lead to over-rotation and under-rotation trends, respectively, during the active stages of OR in eye discs, suggesting that Abl activity in the R3/R4 pair negatively regulates OR.

### Notch recruits dAbl to apical junctional membrane complexes in R4

Fz/PCP signaling appears to be dispensable for the differential apical dAbl localization between R3 and R4, as the pattern is maintained in core PCP mutant ommatidia. Yet, dAbl does synergize with Fmi, when co-over-expressed in the R3/R4 pair, in an OR-specific manner. This OR-associated functional interaction further suggests that dAbl activity is important in R4 at the time of PCP signaling. In other words, dAbl and PCP signaling components are likely to co-operate in the R4 cell to govern the OR process. Our results identify the Notch receptor and signaling in R4 as critical for apical dAbl localization. Notch over-activation within the R3/R4 pair (via expression of stable mutant isoforms of the receptor) induces apical junctional dAbl localization in both cells of the pair. In contrast, expression of a Delta-binding deficient version of Notch (*N^del10-12^*) in R3/R4 pairs, and thus interference with ligand induced Notch activation, frequently leads to a loss of apical dAbl in R4 cells. Similarly, reduction of Notch levels in R3/R4 cells (via RNAi mediated knock-down) also causes a marked decrease in apical dAbl levels in R4. As Notch-dependent transcription is still active in these backgrounds, the combination of these results suggests that the effect of Notch on dAbl apical localization is rather direct, and not via a secondary mechanism through transcriptional regulation. This conclusion is corroborated by the equivalent experiments in salivary gland cells.

One possibility is that the Notch receptor on the membrane physically interacts with dAbl and recruits it to the membrane. Several lines of our data support this hypothesis. In salivary glands, Notch overexpression augments junctional dAbl localization, whereas the total dAbl expression levels are unaffected. In addition, dAbl is co-immunoprecipitates with Notch in third larval instar eye disc extracts, arguing that a membrane associated, direct Notch-Abl interaction mechanism exists *in vivo*, independently of nuclear Notch signaling activity. Notably, both full length and cleavage products of Notch appears to interact with dAbl as well as Arm/β-catenin, suggesting that full length Notch at the membrane is likely involved in the regulation of adherens junction dynamics. Strikingly, the *sev*>*Abl* GOF rotation phenotype is markedly suppressed upon removal of one copy of *Notch*, further supporting the idea that a functional N-Abl signaling module in the apical domain of R4 regulates OR.

### Abl kinase and non-canonical Notch signaling

In *Drosophila*, dAbl has been suggested to act downstream of Notch in the context of axonal pathfinding in embryos. Compelling genetic evidence suggests that a non-canonical Notch signaling branch, which does not entail nuclear Notch activity, instructs axonal pathfinding (Le Gall et al., 2008) and axon-guidance-specific genetic interactions between *dAbl* and *Notch* argue that a non-canonical Notch signaling pathway via dAbl may be at work in this context (Giniger, 1998). Our results are in accordance with these observations and provide further evidence for a potential non-canonical Notch-Abl signaling module to regulate morphogenesis. Recently, a non-canonical Notch pathway has been reported in the regulation of adherens junction organization during human vascular barrier formation (Polacheck et al., 2017). According to this study, the transmembrane domain of Notch complexes with the tyrosine phosphatase LAR, vascular endothelial cadherin, and Rac1 GEF Trio to confer barrier function in human engineered microvessels. The function of the Notch transmembrane domain requires the cleavage of the Notch extracellular and intracellular domains in this context (Polacheck et al., 2017). Our data on the OR process indicate that dAbl apical recruitment in R4 similarly requires Notch activation by Delta. Whether the transmembrane domain of Notch is an essential component of dAbl recruitment and/or regulation remains to be confirmed. However, the observation that dAbl can be co-immunoprecipitated with both the full length as well as the intracellular domain of Notch argues against this idea. In summary, there is a growing body of evidence that Notch utilizes alternative downstream signaling events to regulate cellular morphogenesis and organization, besides its canonical transcriptional target gene regulation (Giniger, 1998; Le Gall et al., 2008; Polacheck et al., 2017; Song and Giniger, 2011).

### Abl kinase regulation of adherens junctions

Our data further demonstrate that Abl affects junctional N-cad levels upon over-expression in the R3/R4 pair. Remarkably, the under-rotation phenotype associated with the dAbl GOF genotype (*sev*>*Abl*) can be suppressed by co-(over)expression of Arm/β-catenin, suggesting that the input from dAbl into adherens junctions is important for the regulation of OR. dAbl is involved in the regulation of multi-cellular reorganization in the context of *Drosophila* germ band elongation through Arm/β-catenin phosphorylation on Tyr667, thus increasing adherens junction turnover to promote convergent extension cell movements (Tamada et al., 2012). dAbl may similarly be involved in regulating Arm/β-catenin dynamics during the OR cell rearrangement process. We have not detected apparent differences in F-actin staining pattern upon dAbl GOF in the R3/R4 cell pair, however, based on previous studies on Abl kinases, it is likely that the actomyosin network is also affected, either directly or indirectly through the adherens junction effect. Further experiments will be needed to test this hypothesis.

### Multiple Notch regulated inputs into the OR process

We have previously reported that Notch signaling in R3/R4 pairs is critical to coordinate OR via Notch-signaling feeding into its transcriptional regulatory mechanism(s) (Koca et al., 2019). Our studies suggested that Notch signaling in R4 directly promotes the transcription of *aos*, encoding an inhibitory ligand to EGFR, and thus fine-tunes the critical EGFR signaling activity (Gaengel and Mlodzik, 2003) within the OR process (Koca et al., 2019). Here, we show that in addition to its interaction with the EGFR signaling pathway, Notch signaling regulates OR via its apical junctional recruitment of dAbl in R4, linking Notch activity here to the non-canonical Abl-mediated Notch pathway and associated local cellular processes. Our data suggest a rather direct role for Notch signaling in regulating OR through the Abl kinase, which then modulates cadherin-based junctional dynamics. Involvement of Notch signaling in cellular morphogenesis has been suggested in various contexts, including *Drosophila* oogenesis and neuronal pathfinding, zebrafish sensory organ development and human vascular barrier formation among others (Cheng et al., 2018; Dobens and Raftery, 2000; Giniger, 1998; Kozlovskaja-Gumbriene et al., 2017; Le Gall et al., 2008; Pang et al., 2021; Polacheck et al., 2017; Song and Giniger, 2011). Besides the reported Notch-signaling mediated transcriptional inputs into adhesion and cytoskeletal dynamics (Grammont, 2007; Kozlovskaja-Gumbriene et al., 2017; Pang et al., 2021), a direct link from the Notch receptor to cell adhesion has also been revealed (Polacheck et al., 2017). Our work here also suggests a direct input from Notch signaling into the cell adhesion factors. Hence, the multifaceted involvement of Notch signaling in cellular (re)organization and morphogenesis is becoming increasingly clear.

Many regulators of the OR process show conservation across critical developmental processes in vertebrates. The reported role of Notch signaling in OR suggests a potential involvement for Notch in PCP-mediated morphogenetic events in vertebrates, which has not been reported thus far. Similarly, Abl kinase may have role in such processes in its interaction with PCP and Notch signaling pathways. Strikingly, the mouse *abl*−/− *arg*−/− double mutants exhibit defects in neurulation and delays in neural tube closure (Koleske et al., 1998), a process generally requiring PCP-regulated features (Butler and Wallingford, 2017; Davey and Moens, 2017; Humphries and Mlodzik, 2018). Future studies will be needed to provide insight into the mechanistic details of how Notch and Abl cooperate in mediating these distinct morphogenetic processes. As Notch signaling and Abl kinase activity have also both long been implicated in cancer metastasis, such studies also hold promise for better understanding of disease and hence future therapeutic applications.

## Materials and Methods

### Fly strains and genetics

Flies were raised on standard medium and maintained at 25°C unless otherwise stated.

*N^1-2155^, N^del10-12^* and *UAS-Abl* were from the Giniger lab.

*UAS-Arm* was a gift from Jennifer Zallen.

*pk^pk-sple13^stbm^6^ FRT42*/CyO *sev-Gal4*/TM3 and *mδ0.5-Gal4 FRT40/SM3:TM6b* were from Mlodzik lab stocks.

*w^1118^*, *N^55e11^/FM6b, abl^2^/TM6b, Notch^RNAi^* (BL31383) and *Abl^RNAi^* (BL35327) lines were ordered from Bloomington Drosophila Stock Center.

*mδ0.5*>*N^RNAi BL31383^* (*mδ0.5-Gal4*/+; *UAS-N^RNAi BL31383^*/+) were obtained at 25°C.

*mδ0.5*>*N^1-2155^* (*mδ0.5-Gal4*/+; *UAS-N^1-2155^*/+) were obtained at 25°C.

*mδ0.5*>*N^del10-12^ (mδ0.5-Gal4/+; UAS-N^del10-12^/+)* were obtained at 29°C.

*mδ0.5*>*Abl^RNAi BL35327^ (mδ0.5-Gal4/+; UAS-Abl^RNAi BL35327^ /+)* were obtained at 29°C.

*mδ0.5*>*Abl^RNAi VDRC110186^ (mδ0.5-Gal4/+; UAS-Abl^RNAi VDRC110186^/+)* were obtained at 29°C.

*mδ0.5*>*Abl (mδ0.5-Gal4, UAS-Abl*/+) were obtained at 29°C.

*sev*>*Abl (sev-Gal4*/+; *UAS-Abl*/+) were obtained at 18°C.

Control eye disc stainings were done in *mδ0.5-Gal4 FRT40*/+ backgrounds.

Genetic interactions were tested at 18°C between *sev-Gal4/+; UAS-Abl/+* and the heterozygosity of the respective transgenic flies.

*abl^2^* mutants were generated with the following genotype:

*eyFLP/+; abl^2^ FRT80/ubiGFP FRT80*

*pk^pk-sple13^stbm^6^* double mutants were generated with the following genotype:

*eyFLP/+; pk^pk-sple13^stbm^6^ FRT42/ubiGFP FRT42*

*mδ0.5>UAS-Abl* clones were obtained at 29°C by employing FLP/FRT mediated mitotic recombination with the following genotypes:

*eyFLP/+; mδ0.5-Gal4 FRT40/ubiGFP FRT40; UAS-Abl/+*

### Immunohistochemistry and Histology

Adult eye sectioning was performed as previously described (Jenny, 2011).

Third larval instar eye discs were dissected in ice-cold PBS and fixed in PBT (PBS+0.1% Triton-X)-4% formaldehyde for 12 minutes at room temperature. For immunohistochemistry, following primary antibodies were used: rabbit anti-Abl (1:125, Mlodzik Lab), rat anti-DE-cad (1:20, DSHB), mouse anti-Fmi (1:10, DSHB), rat anti-DN-cad (1:20, DSHB), mouse anti-Arm (1:20, DSHB), chicken anti-GFP (1:1000, Aves Labs) and rhodamine phalloidin (1:1000, Invitrogen). Secondary antibodies were obtained from Jackson Laboratories. Images were acquired by using Leica SP5 DMI microscope.

### Biochemical interaction assays

For *in vitro* GST pull downs, the recombinant proteins GST-Notch ICD, GST-Su(H) and MBP-Abl were expressed in bacteria and purified. For the amplification of Notch ICD, Abl and Su(H) fragments, the following primers were used:

Notch forward primer: 5’ CATGCCCGGGACAAAGAAAGCGGGCACATG

Notch reverse primer: 5’ CTAGGCGGCCGCTCAAATGTAGATGGCCTC

Abl forward primer: 5’ CATGTCTAGAGGGGCTCAGCAGGGCAAGG

Abl reverse primer: 5’ CATGGTCGACTTACCTGTTAAGCGCATTGG

Su(H) forward primer: 5’ ATGAGAATTCATGTGTGATTAGTCGTGAATC

Su(H) reverse primer: 5’ ATGAGTCGACTCATTTAGATCTTTGGAAATTCAT

Amplified fragments were cloned into pGEX-4T1 vector for GST tagging and pMAL-c2X vector for MBD tagging. Bacteria lysates were prepared as described (Frangioni and Neel, 1993). Glutathion Sepharose 4B beads (GE Healthcare) with GST or with GST-fusion proteins were incubated with lysates containing MBP-Abl O/N at 4°C. After that, the samples were washed 3 times with pull-down buffer (20 mM Tris pH 7.6, 150 mM NaCl, 0.5 mM EDTA, 10% glycerol, 0.1% Triton X 100, 1mM DTT, and protease inhibitor cocktail). Pull-down samples were resuspended in SDS sample buffer, boiled for 5 min and the proteins were separated by SDS PAGE. For Western blot, the nitrocellulose membrane was incubated with mouse-anti-GST and rabbit-anti-MBP antibody. Proteins were detected by enhanced chemiluminescence (Millipore).

For *in vivo* co-immunoprecipitations eye imaginal discs were dissected from third instar larvae. Lysates from 30 wt eye imaginal discs (1 mg of total protein) was precleared by incubating with protein A-sepharose beads (Thermo Scientific) for 1h at 4 °C followed by centrifugation. A-sepharose beads were immunoprecipitated with Abl or Notch antibody and the lysates at 4 °C O/N. Immunoprecipitates were captured by protein A-sepharose at 4 °C in IP buffer (20 mM 4-(2-hydroxyethyl)-1-piperazineethanesulfonic acid (HEPES) pH 7.5, 100 mM NaCl, 0.05% Triton X100, 1 mM EDTA, 5 mM DTT, 10 mM NaVO4, 10% glycerol and protease inhibitor cocktail). After several washes with IP buffer, immunoprecipitates were resuspended in SDS sample buffer; beads were boiled for 5 min, and proteins were resolved on SDS-PAGE. For western blotting, proteins were transferred onto nitrocellulose membrane, blocked in 5% skim milk (Labscientific) and incubated with primary rabbit anti-Abl or mouse anti-NotchICD, and secondary anti-rabbit or anti-mouse antibody. Protein was detected by enhanced chemiluminescence (Millipore).

### Quantitative Analysis of Adult Eye Sections

The orientation of each ommatidium was marked based on the trapezoidal organization of the R-cells (see Figure 1A). A linear equator has been drawn along the boundary where two chiral forms meet. Clockwise and counter-clockwise angles from the equator to each ommatidia were measured for the black and red chiral forms respectively (see Figure 1A, C). Measurements were done by using ImageJ (National Institute of Health). The absolute values of measured angles from 3-4 independent eye sections for each genotype were pooled (300<n<500) and plotted in a histogram by using MATLAB. The angles were binned into 20° intervals between 0-180° and they were plotted in probability ratios from 0 to 1. For statistical analyses, the angles (α) were binned into 3 categories (α<60, 60<α<120, 120<α) for individual genotypes and chi-square test was performed.

### Quantitative Analysis of Eye Discs

The orientation of each ommatidium was marked perpendicular to the plane of R2/R5 cells (See Figure 1B,2C-D). A linear equator was drawn perpendicular to the MF at the dorsoventral midline. Clockwise and counter-clockwise angles from the equator to each ommatidia were measured for the dorsal and ventral halves respectively. To avoid a potential bias due to the developmental delay in rotation from equator to the poles, the measurements were limited to the first 8 ommatidia from the equator for each row. Measurements were done by using ImageJ (National Institute of Health). The absolute values of measured angles from 5-8 independent eye discs (30<n<70 per row) were pooled and violin plotted in PRISM. For statistical analyses, the angles (α) from individual rows were binned into 5 categories (α<40, 40< α<50, 50< α<60, 60< α<70 and α>70) for each genotype and chi-square test was performed.

For Abl quantifications in control and Notch-perturbed backgrounds, apical confocal stacks were maximum projected and individual cell intensities were measured in R4 and R2 cells between rows 9-11 by using ImageJ. Intensities in R4 cells were normalized to their R2 neighbors within each ommatidia. Measurements from 3 discs were pooled (45<n<60 per genotype) and plotted. For statistical analyses, the normalized intensity measurements (*ι*) were binned into 2 categories (*ι*<1, *ι*>1) for each genotype and chi-square test was performed.

For DN-cad quantifications in *mδ0.5*>*Abl* mosaic eye discs, apical confocal stacks were maximum projected and integrated DN-cad intensities were measured in ImageJ. Within GFP^+^ and GFP^−^ tissue, DN-cad integrated intensity within the R3/R4 boundary was measured for each ommatidia between rows 5-11 and plotted. Measurements from 3 discs were pooled (35<n<70 per genotype) and plotted. For statistical analyses, the normalized intensity measurements (*ι*) were binned into 2 categories (*ι*<10, *ι*>10) for each genotype and chi-square test was performed.

For Arm/β-catenin quantifications in *mδ0.5*>*Abl* mosaic eye discs, apical confocal stacks were maximum projected and Arm intensities between R3/R4 borders were measured in ImageJ. Within GFP^+^ and GFP^−^ tissue, Arm intensity at 3 different points along the R3/R4 boundary was measured for each ommatidia between rows 5-11 and the mean intensity for each ommatidia was plotted. Measurements from 3 discs were pooled (35<n<70 per genotype) and plotted. For statistical analyses, mean intensity measurements (*ι*) were binned into 2 categories (*ι*<0.5, *ι*>0.5) for each genotype and chi-square test was performed.

### Quantitative Analysis of Salivary Glands

Salivary gland staining images were processed using ImageJ. For apical Notch and Abl quantifications, apical confocal stacks were maximum projected, apical junctional intensities were measured at multiple points and normalized to the corresponding DE-cad intensities followed by averaging for each salivary gland. Mean measurements were plotted for 12 salivary glands. For statistical analyses, Student’s two tailed t-test was performed on normalized mean intensity measurements.

Western blot images were processed using ImageJ. For relative protein expression quantifications of Notch and Abl, scans of developed films were converted to 8-bit images to perform uncalibrated optical density. Each band was individually selected and circumscribed with the rectangular ROI selection, followed by quantification of peak area of obtained histograms. Data were acquired with arbitrary area values. The intensity of protein bands was normalized to expression of Tubulin. For statistical analyses, Student’s two tailed t-test was performed on normalized intensity measurements.

## Acknowledgements

We thank the Bloomington Drosophila Research Center and Jennifer Zallen for fly strains and reagents. We are grateful to all Mlodzik lab members, past and present, members for helpful input and discussions; and Robert Krauss, Cathie Pfleger, Timothy Blenkinsop and Jennifer Zallen for helpful comments and suggestions on the manuscript. Confocal laser scanning microscopy was performed at the ISMMS-Microscopy Core Facility supported by the Tisch Cancer Institute grant P30 CA196521 from the NCI. This work was supported by NIH/NEI grant R01 EY13256 to M.M.

## Author contributions

Y.K. and M.M. conceived and designed the study. Y.K. performed immunohistochemistry, histology, genetic, and functional experiments and analyzed the data. L.V. performed biochemical interaction and salivary gland experiments, and analyzed the data. J.S. performed the genetic interaction experiments between Abl and Notch. E.G. provided conceptual input and the Notch truncation constructs, which he tested. Y.K. and M.M. wrote the paper with input from all authors. M.M. acquired the research funds for the project.

## Competing interests

The authors declare no competing interests.

## Notes

### Competing Interest Statement

The authors have declared no competing interest.

## References

Adler, P.N. (2012). The frizzled/stan pathway and planar cell polarity in the Drosophila wing. Curr Top Dev Biol 101, 1–31.

Baum, B., and Perrimon, N. (2001). Spatial control of the actin cytoskeleton in Drosophila epithelial cells. Nat Cell Biol 3, 883–890.

Blair, S.S. (1999). Eye development: Notch lends a handedness. Curr Biol 9, R356–360.

Bradley, W.D., and Koleske, A.J. (2009). Regulation of cell migration and morphogenesis by Abl-family kinases: emerging mechanisms and physiological contexts. J Cell Sci 122, 3441–3454.

Brown, K.E., and Freeman, M. (2003). Egfr signalling defines a protective function for ommatidial orientation in the Drosophila eye. Development 130, 5401–5412.

Butler, M.T., and Wallingford, J.B. (2017). Planar cell polarity in development and disease. Nat Rev Mol Cell Biol 18, 375–388.

Cagan, R.L., and Ready, D.F. (1989). The emergence of order in the Drosophila pupal retina. Dev Biol 136, 346–362.

Cheng, D., Yan, X., Qiu, G., Zhang, J., Wang, H., Feng, T., Tian, Y., Xu, H., Wang, M., He, W., et al. (2018). Contraction of basal filopodia controls periodic feather branching via Notch and FGF signaling. Nat Commun 9, 1345.

Choi, K.W., and Benzer, S. (1994). Rotation of photoreceptor clusters in the developing Drosophila eye requires the nemo gene. Cell 78, 125–136.

Chou, Y.H., and Chien, C.T. (2002). Scabrous controls ommatidial rotation in the Drosophila compound eye. Dev Cell 3, 839–850.

Cooper, M.T., and Bray, S.J. (1999). Frizzled regulation of Notch signalling polarizes cell fate in the Drosophila eye. Nature 397, 526–530.

Das, G., Reynolds-Kenneally, J., and Mlodzik, M. (2002). The atypical cadherin Flamingo links Frizzled and Notch signaling in planar polarity establishment in the Drosophila eye. Dev Cell 2, 655–666.

Davey, C.F., and Moens, C.B. (2017). Planar cell polarity in moving cells: think globally, act locally. Development 144, 187–200.

Devenport, D. (2016). Tissue morphodynamics: Translating planar polarity cues into polarized cell behaviors. Semin Cell Dev Biol 55, 99–110.

Dobens, L.L., and Raftery, L.A. (2000). Integration of epithelial patterning and morphogenesis in Drosophila ovarian follicle cells. Dev Dyn 218, 80–93.

Fanto, M., and Mlodzik, M. (1999). Asymmetric Notch activation specifies photoreceptors R3 and R4 and planar polarity in the Drosophila eye. Nature 397, 523–526.

Fiehler, R.W., and Wolff, T. (2007). Drosophila Myosin II, Zipper, is essential for ommatidial rotation. Dev Biol 310, 348–362.

Fiehler, R.W., and Wolff, T. (2008). Nemo is required in a subset of photoreceptors to regulate the speed of ommatidial rotation. Dev Biol 313, 533–544.

Fox, D.T., and Peifer, M. (2007). Abelson kinase (Abl) and RhoGEF2 regulate actin organization during cell constriction in Drosophila. Development 134, 567–578.

Frangioni, J.V., and Neel, B.G. (1993). Solubilization and purification of enzymatically active glutathione S-transferase (pGEX) fusion proteins. Anal Biochem 210, 179–187.

Gaengel, K., and Mlodzik, M. (2003). Egfr signaling regulates ommatidial rotation and cell motility in the Drosophila eye via MAPK/Pnt signaling and the Ras effector Canoe/AF6. Development 130, 5413–5423.

Giniger, E. (1998). A role for Abl in Notch signaling. Neuron 20, 667–681.

Goodrich, L.V., and Strutt, D. (2011). Principles of planar polarity in animal development. Development 138, 1877–1892.

Grammont, M. (2007). Adherens junction remodeling by the Notch pathway in Drosophila melanogaster oogenesis. J Cell Biol 177, 139–150.

Grevengoed, E.E., Fox, D.T., Gates, J., and Peifer, M. (2003). Balancing different types of actin polymerization at distinct sites: roles for Abelson kinase and Enabled. J Cell Biol 163, 1267–1279.

Grevengoed, E.E., Loureiro, J.J., Jesse, T.L., and Peifer, M. (2001). Abelson kinase regulates epithelial morphogenesis in Drosophila. J Cell Biol 155, 1185–1198.

Hantschel, O., Nagar, B., Guettler, S., Kretzschmar, J., Dorey, K., Kuriyan, J., and Superti-Furga, G. (2003). A myristoyl/phosphotyrosine switch regulates c-Abl. Cell 112, 845–857.

Hernández, S.E., Krishnaswami, M., Miller, A.L., and Koleske, A.J. (2004). How do Abl family kinases regulate cell shape and movement? Trends in Cell Biology 14, 36–44.

Humphries, A.C., and Mlodzik, M. (2018). From instruction to output: Wnt/PCP signaling in development and cancer. Curr Opin Cell Biol 51, 110–116.

Jenny, A. (2010). Planar cell polarity signaling in the Drosophila eye. Curr Top Dev Biol 93, 189–227.

Jenny, A. (2011). Preparation of adult Drosophila eyes for thin sectioning and microscopic analysis. J Vis Exp.

Jenny, A., Darken, R.S., Wilson, P.A., and Mlodzik, M. (2003). Prickle and Strabismus form a functional complex to generate a correct axis during planar cell polarity signaling. EMBO J 22, 4409–4420.

Jodoin, J.N., and Martin, A.C. (2016). Abl suppresses cell extrusion and intercalation during epithelium folding. Mol Biol Cell 27, 2822–2832.

Kannan, R., Song, J.K., Karpova, T., Clarke, A., Shivalkar, M., Wang, B., Kotlyanskaya, L., Kuzina, I., Gu, Q., and Giniger, E. (2017). The Abl pathway bifurcates to balance Enabled and Rac signaling in axon patterning in Drosophila. Development 144, 487–498.

Koca, Y., Housden, B.E., Gault, W.J., Bray, S.J., and Mlodzik, M. (2019). Notch signaling coordinates ommatidial rotation in the Drosophila eye via transcriptional regulation of the EGF-Receptor ligand Argos. Sci Rep 9, 18628.

Koleske, A.J., Gifford, A.M., Scott, M.L., Nee, M., Bronson, R.T., Miczek, K.A., and Baltimore, D. (1998). Essential roles for the Abl and Arg tyrosine kinases in neurulation. Neuron 21, 1259–1272.

Kozlovskaja-Gumbriene, A., Yi, R., Alexander, R., Aman, A., Jiskra, R., Nagelberg, D., Knaut, H., McClain, M., and Piotrowski, T. (2017). Proliferation-independent regulation of organ size by Fgf/Notch signaling. Elife 6.

Le Gall, M., De Mattei, C., and Giniger, E. (2008). Molecular separation of two signaling pathways for the receptor, Notch. Dev Biol 313, 556–567.

Mirkovic, I., Gault, W.J., Rahnama, M., Jenny, A., Gaengel, K., Bessette, D., Gottardi, C.J., Verheyen, E.M., and Mlodzik, M. (2011). Nemo kinase phosphorylates beta-catenin to promote ommatidial rotation and connects core PCP factors to E-cadherin-beta-catenin. Nat Struct Mol Biol 18, 665–672.

Mirkovic, I., and Mlodzik, M. (2006). Cooperative activities of drosophila DE-cadherin and DN-cadherin regulate the cell motility process of ommatidial rotation. Development 133, 3283–3293.

Mlodzik, M. (1999). Planar polarity in the Drosophila eye: a multifaceted view of signaling specificity and cross-talk. EMBO J 18, 6873–6879.

Nagar, B., Hantschel, O., Young, M.A., Scheffzek, K., Veach, D., Bornmann, W., Clarkson, B., Superti-Furga, G., and Kuriyan, J. (2003). Structural basis for the autoinhibition of c-Abl tyrosine kinase. Cell 112, 859–871.

Pang, J., Le, L., Zhou, Y., Tu, R., Hou, Q., Tsuchiya, D., Thomas, N., Wang, Y., Yu, Z., Alexander, R., et al. (2021). NOTCH Signaling Controls Ciliary Body Morphogenesis and Secretion by Directly Regulating Nectin Protein Expression. Cell Rep 34, 108603.

Peng, Y., and Axelrod, J.D. (2012). Asymmetric protein localization in planar cell polarity: mechanisms, puzzles, and challenges. Curr Top Dev Biol 101, 33–53.

Polacheck, W.J., Kutys, M.L., Yang, J., Eyckmans, J., Wu, Y., Vasavada, H., Hirschi, K.K., and Chen, C.S. (2017). A non-canonical Notch complex regulates adherens junctions and vascular barrier function. Nature 552, 258–262.

Rebay, I., Fleming, R.J., Fehon, R.G., Cherbas, L., Cherbas, P., and Artavanis-Tsakonas, S. (1991). Specific EGF repeats of Notch mediate interactions with Delta and Serrate: implications for Notch as a multifunctional receptor. Cell 67, 687–699.

Roignant, J.Y., and Treisman, J.E. (2009). Pattern formation in the Drosophila eye disc. Int J Dev Biol 53, 795–804.

Singh, J., Yanfeng, W.A., Grumolato, L., Aaronson, S.A., and Mlodzik, M. (2010). Abelson family kinases regulate Frizzled planar cell polarity signaling via Dsh phosphorylation. Genes Dev 24, 2157–2168.

Song, J.K., and Giniger, E. (2011). Noncanonical Notch function in motor axon guidance is mediated by Rac GTPase and the GEF1 domain of Trio. Dev Dyn 240, 324–332.

Song, J.K., Kannan, R., Merdes, G., Singh, J., Mlodzik, M., and Giniger, E. (2010). Disabled is a bona fide component of the Abl signaling network. Development 137, 3719–3727.

Strutt, H., and Strutt, D. (1999). Polarity determination in the Drosophila eye. Curr Opin Genet Dev 9, 442–446.

Tamada, M., Farrell, D.L., and Zallen, J.A. (2012). Abl regulates planar polarized junctional dynamics through beta-catenin tyrosine phosphorylation. Dev Cell 22, 309–319.

Tomlinson, A., and Ready, D.F. (1987). Neuronal differentiation in Drosophila ommatidium. Dev Biol 120, 366–376.

Tomlinson, A., and Struhl, G. (1999). Decoding vectorial information from a gradient: sequential roles of the receptors Frizzled and Notch in establishing planar polarity in the Drosophila eye. Development 126, 5725–5738.

Wang, J.Y. (2014). The capable ABL: what is its biological function? Mol Cell Biol 34, 1188–1197.

Weber, U., Paricio, N., and Mlodzik, M. (2000). Jun mediates Frizzled-induced R3/R4 cell fate distinction and planar polarity determination in the Drosophila eye. Development 127, 3619–3629.

Weber, U., Pataki, C., Mihaly, J., and Mlodzik, M. (2008). Combinatorial signaling by the Frizzled/PCP and Egfr pathways during planar cell polarity establishment in the Drosophila eye. Dev Biol 316, 110–123.

Winter, C.G., Wang, B., Ballew, A., Royou, A., Karess, R., Axelrod, J.D., and Luo, L. (2001). Drosophila Rho-associated kinase (Drok) links Frizzled-mediated planar cell polarity signaling to the actin cytoskeleton. Cell 105, 81–91.

Wolff, T., and Ready, D.F. (1991). The beginning of pattern formation in the Drosophila compound eye: the morphogenetic furrow and the second mitotic wave. Development 113, 841–850.

Wolff, T., and Rubin, G.M. (1998). Strabismus, a novel gene that regulates tissue polarity and cell fate decisions in Drosophila. Development 125, 1149–1159.

Wu, J., Klein, T.J., and Mlodzik, M. (2004). Subcellular localization of frizzled receptors, mediated by their cytoplasmic tails, regulates signaling pathway specificity. PLoS Biol 2, E158.

Wu, J., and Mlodzik, M. (2009). A quest for the mechanism regulating global planar cell polarity of tissues. Trends Cell Biol 19, 295–305.

Xiong, W., and Rebay, I. (2011). Abelson tyrosine kinase is required for Drosophila photoreceptor morphogenesis and retinal epithelial patterning. Dev Dyn 240, 1745–1755.

Zheng, L., Zhang, J., and Carthew, R.W. (1995). frizzled regulates mirror-symmetric pattern formation in the Drosophila eye. Development 121, 3045–3055.

