## Supplemental Figures for "Notch-dependent Abl signaling regulates cell motility during ommatidial rotation in *Drosophila*"

### Supplemental Data

### Supplemental Figures

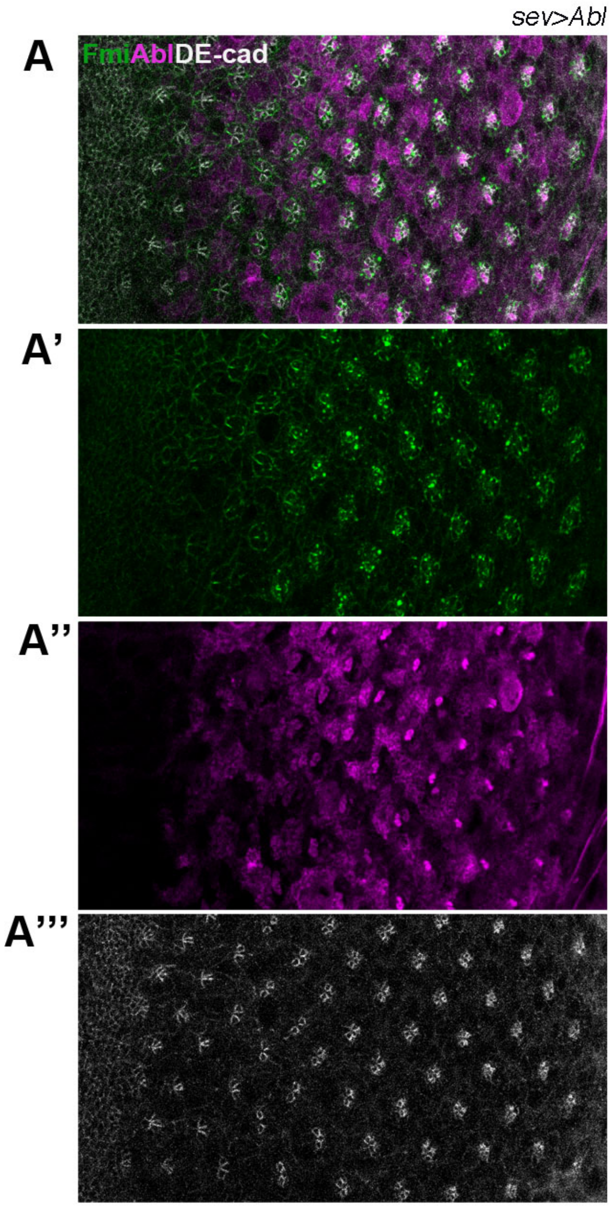


**Figure S2 (to Figure 2)**. Third instar larval eye imaginal discs stained for Fmi (green), Abl (magenta) and DE-cad (gray) in *sev>Abl* background. Note that sev-driven Abl is overexpressed in many cells behind the furrow (not just the *sev*-competence group cells of R3/R4, R1/R6, R7 and the cone cells). This is likely due to the combination of the *sev*-enhancer and UAS-transgene chromosomal position insertions.

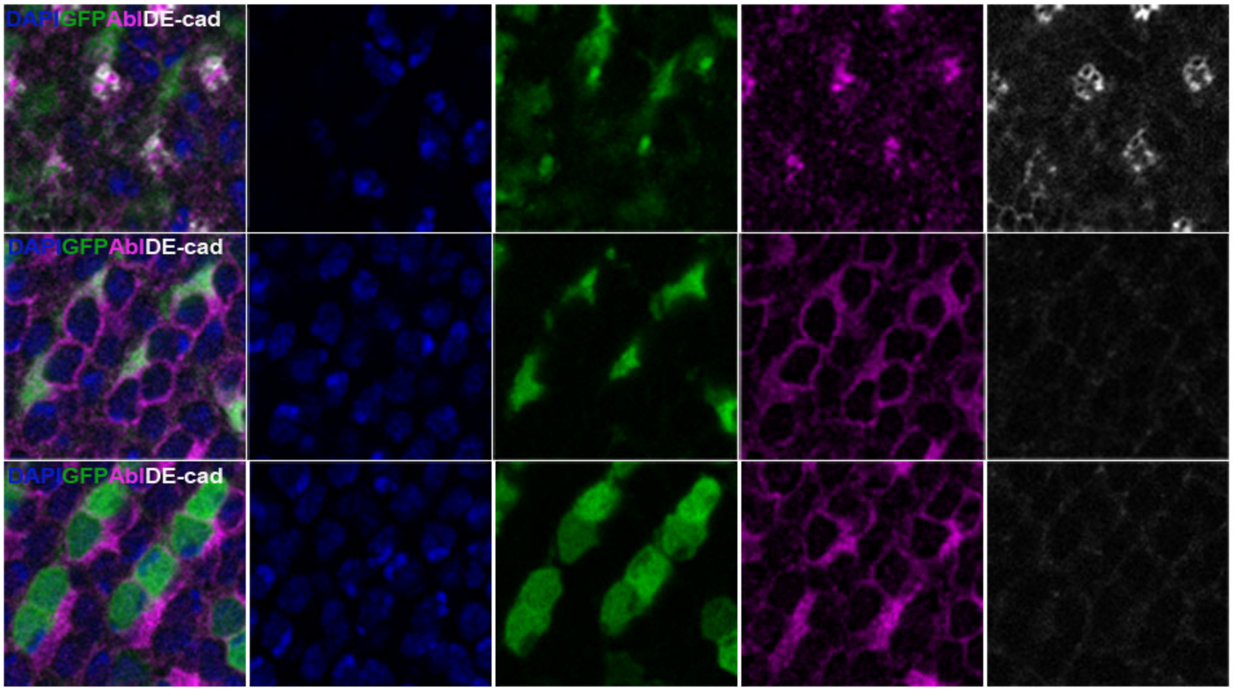


**Figure S3 (to Figure 3)**. Third instar larval eye imaginal discs of *mδ0.5>GFP* genotype, stained for GFP (green), Abl (magenta), DE-cad (gray) and DAPI (blue). The slices were depicted for three planes from apical (top row) to basal (bottom row). GFP is expressed only in R3/R4 cells (high in R4, low in R3). Note that Abl staining do not exhibit difference between R3/R4 at the subapical sections, whereas it is enriched only in R4 at the most apical slice.

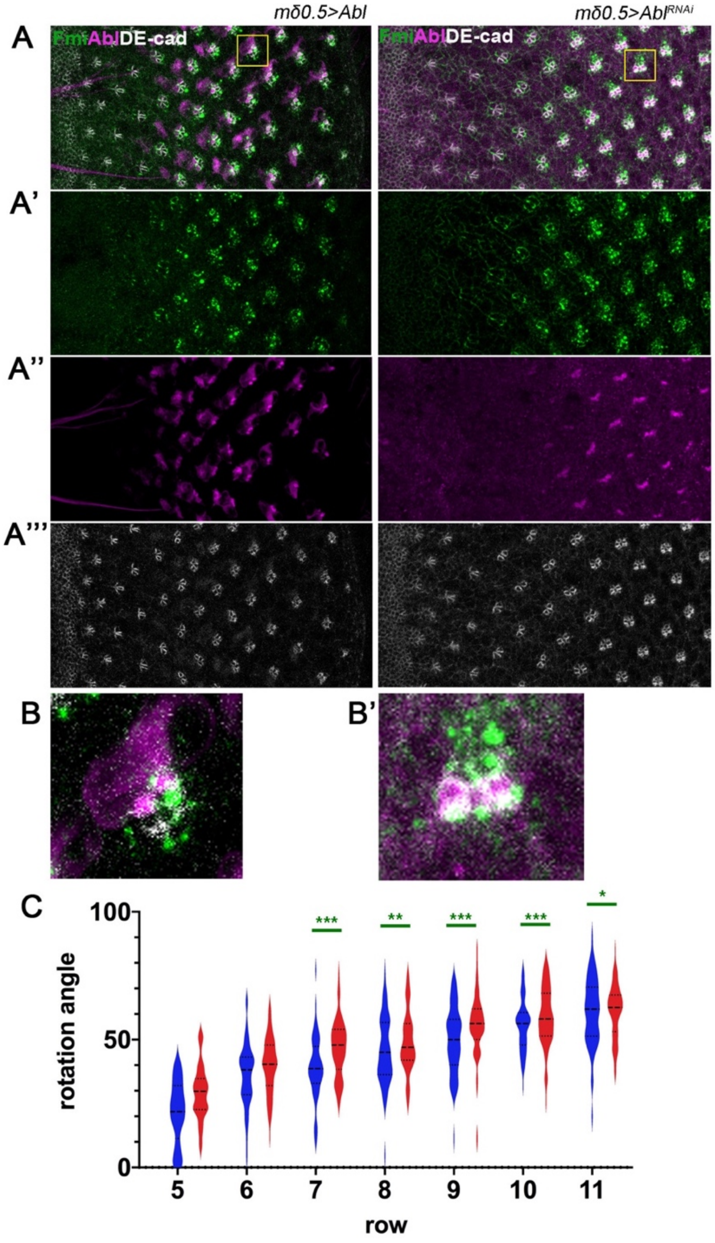


**Figure S4 (to Figure 4)**. (A-A''') Third instar larval eye imaginal discs stained for Fmi (green), Abl (magenta) and DE-cad (gray) in *mδ0.5>Abl* (left panel column), and *mδ0.5>Abl*$^{RNAi\#2}$ backgrounds (right panel column). (B-B'): Individual clusters for each genotype (from yellow boxes in A) shown at high magnification. Note that Abl is expressed in both R3 and R4 cells in *mδ0.5>Abl* (B) background and it is depleted from the R4 apical space in *mδ0.5>Abl* (B') background. (C) Quantification of rotation angles observed in individual preclusters in rows 5–11, plotted for *wt* (blue) and *mδ0.5>Abl*$^{RNAi\#2}$ (red).

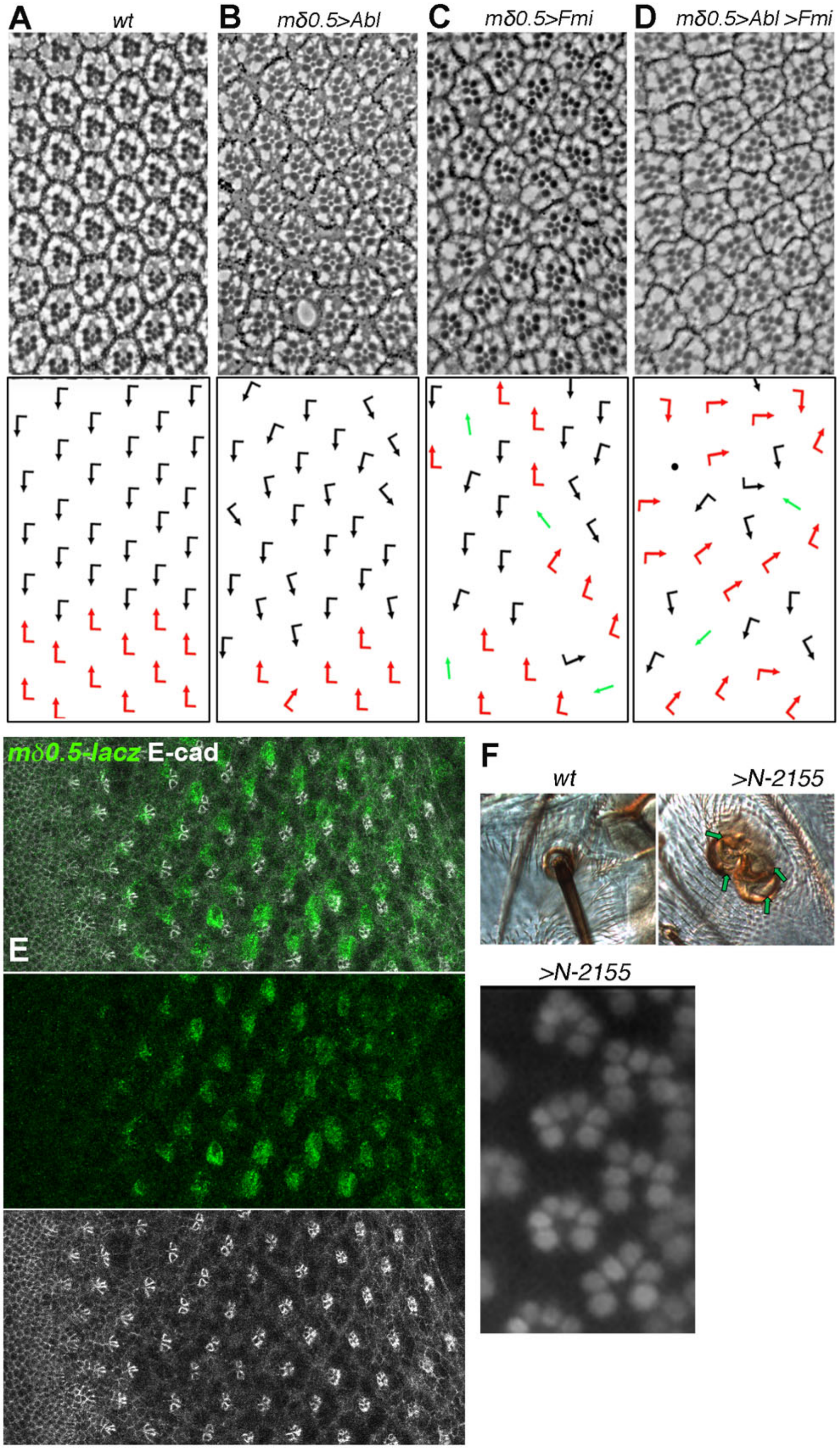


**Figure S5 (to Figure 5)**. Adult eye sections with ommatidial orientation schematics and orientation angle histograms of eyes of the genotypes indicated: (A) *wt*, (B) *mδ0.5>Abl*, (C) *mδ0.5>Fmi* (D)

*mδ0.5>Abl>Fmi.* Note that ommatidia are misoriented in D to the extent that equator is not distinguishable.

(E) Third instar larval eye imaginal discs of *mδ0.5>*$N^{del10-12}$, *mδ0.5-lacZ* background stained for β-gal (green) and E-cad (white). Note that *mδ0.5-lacZ* is still active in *mδ0.5>*$N^{del10-12}$ background. (F) Expression of the $N^{1-2155}$ truncatied isoform behaves like an activated receptor inducing Notch GOF phenotypes on the thorax (upper panels) and the eye (lower panel). In the SOP group on the thorax GOF Notch (expressed with the *109-68* Gal4 driver) shows a four-socket phenotype (compare to *wt* on left), and in the eye (expressed with *sev-Gal4* or *mδ0.5-Gal4*) it can cause R4/R4 symmetrical ommatidial clusters (left side of panel, compare to clusters with *wt* chiral arrangement on right side of panel).

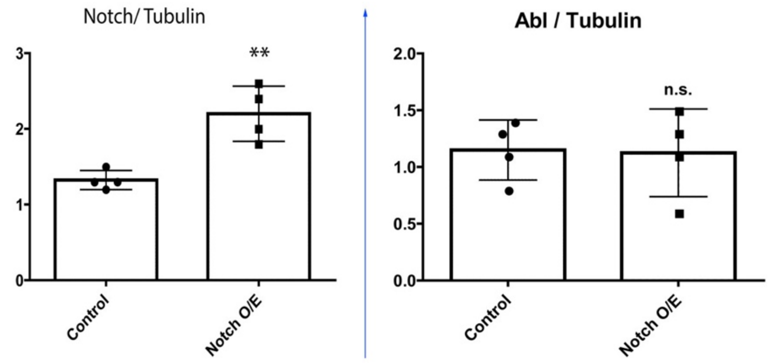


**Figure S6 (to Figure 6)**. Western blot quantifications of Notch and Abl protein levels (total cell extracts) in *wt* and Notch overexpressing salivary glands immunoblotted for Notch and Abl (and normalized to Tubulin). Note that while overall Notch levels increase (due to the Gal4 driven overexpression), total Abl levels are not affected, indicating that Notch causes a specific enrichment of Abl at the junctions (as shown in Fig. 6F-H).

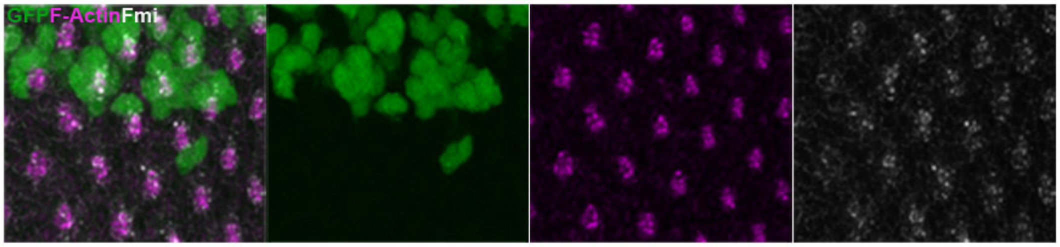


**Figure S7 (to Figure 7)**. *mδ0.5>Abl* mosaic eye disc stained for GFP (green), F-actin (magenta) and Fmi (gray). *mδ0.5>Abl* tissue is marked by the absence of GFP. Note that F-actin distribution in preclusters is not affected upon Abl overexpression.
